# PI-(3,5)P2-mediated oligomerization of the endosomal sodium/proton exchanger NHE9

**DOI:** 10.1101/2023.09.10.557050

**Authors:** Surabhi Kokane, Pascal F. Meier, Ashutosh Gulati, Rei Matsuoka, Tanadet Pipatpolkai, Giuseppe Albano, Lucie Delemotte, Daniel Fuster, David Drew

## Abstract

Na^+^/H^+^ exchangers are found in all cells to regulate intracellular pH, sodium levels and cell volume. Na^+^/H^+^ exchangers are physiological homodimers that operate by an elevator alternating-access mechanism. While the structure of the core ion translocation domain is fairly conserved, the scaffold domain and oligomerization show larger structural variation. The Na^+^/H^+^ exchanger NhaA from *E. coli* has a weak oligomerization interface mediated by a β-hairpin domain and homodimerization was shown to be dependent of the lipid cardiolipin. Organellar Na^+^/H^+^ exchangers NHE6, NHE7 and NHE9 are likewise predicted to contain β-hairpin domains and a recent analysis of *horse* NHE9 indicated that the lipid PIP_2_ binds at the dimerization interface. Despite predicted lipid-mediated oligomerization, their structural validation has been lacking. Here, we report cryo-EM structures of *E. coli* NhaA and *horse* NHE9 with the coordination of cardiolipin and PI(3,5)P_2_ binding at the dimer interface, respectively. Cell based assays confirms that NHE9 is inactive at the plasma membrane and thermal-shift assays, solid-supported membrane (SSM) electrophysiology and MD simulations, corroborates that NHE9 specifically binds the endosomal PI(3,5)P_2_ lipid, which stabilizes the homodimer and enhances activity. Taken together, we propose specific lipids regulate Na^+^/H^+^ exchange activity by stabilizing oligomerization and stimulating Na^+^ binding under lipid-specific cues.

## Introduction

Na^+^/H^+^ exchangers facilitate the exchange of cations (Na^+^/Li^+^/K^+^) and protons (H^+^) across membranes to regulate intracellular pH, sodium levels and cell volume (Pedersen & Counillon, 2019). In bacteria these ancient proteins work to either generate a sodium-motive-force or to alleviate sodium-salt stress (Krulwich *et al*, 2011; Padan, 2008). In mammals, there are 13 different Na^+^/H^+^ exchangers (NHE) belonging to the SLC9A (NHE1-9), SLC9B (NHA1-2) or sperm specific SLC9C families (Fuster & Alexander, 2014; Pedersen & Counillon, 2019). NHEs differ in their substrate preferences, kinetics, subcellular and tissue distribution (Pedersen & Counillon, 2019). Isoforms NHE1-5 are primarily expressed on the plasma membrane and have important physiological roles linked to cellular pH homeostasis (Pedersen & Counillon, 2019). Isoforms NHE6-9 are primary localized to intracellular compartments and work in concert with the V-type ATPase to fine-tune pH in their respective organelles.

Na^+^/H^+^ exchangers are physiological homodimers belonging to the “NhaA-fold”, so-named after the first fold-representative to be determined (Brett *et al*, 2005; Fliegel, 2019; Padan, 2008). The transporter is made up of two distinct domains, a dimerization domain and an ion-transporting (core) domain. The 6-TM core domain undergoes global, elevator-like structural transitions to translocate ions across the membrane against the anchored dimerization domain (Coincon *et al*, 2016; Drew & Boudker, 2016). The cryo-EM structures of SLC9A, SLC9B and SLC9C have been determined and show a similar architecture to bacterial homologue structures with 13 transmembrane (TM) helices (Hunte *et al*, 2005a; Lee *et al*, 2013; Matsuoka *et al*, 2022; Paulino *et al*, 2014; Wohlert *et al*, 2014; Yeo *et al*, 2023). Folds of secondary-active transporters are nearly always made up from a conserved repeat unit that is related by internal structural-symmetry (Forrest, 2015). Interestingly, the structural-inverted repeats in proteins with the NhaA-fold seem to be the only transporter members where the structural-repeat unit varies between members. NhaA is made up of 12 TMs and has a 5-TM structural-inverted repeat, SLC9A and SLC9C are made up of 13-TMs and have a 6-TM repeat, whereas SLC9B has 14-TMs and a 7-TM repeat (Matsuoka *et al*., 2022; Yeo *et al*., 2023). Although the 6-TM core domain of NhaA that translocates the ions across the membrane is conserved, the additional number of helices alters the structure of the dimerization interface. It seems that there has been evolutionary pressure to fine-tune oligomerization in the NhaA-fold for a functional reason.

Previous studies using native MS, MD simulations and thermal-shift assays, identified that NhaA oligomerization is dependent on cardiolipin (Gupta *et al*, 2017; Nji *et al*, 2018; Rimon *et al*, 2019; Winkelmann *et al*, 2022). NhaA has a weak dimer interface mediated by a β-hairpin domain, which may make dimerization and activity more lipid-dependent than of other Na^+^/H^+^ exchangers. Consistent with this reasoning, dimerization in other bacterial Na^+^/H^+^ exchangers lacking β-hairpins and with a larger dimerization interface, were found to be lipid-independent (Gupta *et al*., 2017). It was further shown that NhaA was unable to compensate salt-stress in a *E. coli* strain deficient in cardiolipin (Rimon *et al*., 2019). Since cardiolipin biosynthesis in *E. coli* is increased upon salt-stress and, because NhaA is required to alleviate salt-stress (Padan, 2008; Romantsov *et al*, 2009), it was thus proposed that lipid binding might be a means to regulate *in vivo* NhaA activity. Despite this attractive hypothesis, structural validation of cardiolipin binding to *E. coli* NhaA has been lacking.

In addition to NhaA, native mass-spectrometry and thermal-shift assays have also shown that the endosomal Na^+^/H^+^ exchanger NHE9 (SLC9A9) dimer co-purifies with negatively-charged lipids matching the mass of PIP_2_ (Winklemann *et al*, 2020). NHE9 regulates the luminal pH of late– and recycling endosomes (Kondapalli *et al*, 2015; Kondapalli *et al*, 2014) and mutations in SLC9A9 have been associated with neurological disorders such as familial autism, ADHD and epilepsy (Kondapalli *et al*., 2014; Zhang-James *et al*, 2019). Interestingly, although endosomal NHE9 has a larger dimer interface than NhaA and is similar in size to plasma membrane localised NHE1 (Dong *et al*, 2021) (Fig. 1A), the NHE9 protein has an additional ∼ 60 residue extension of the TM2-TM3 loop, which AlphaFold 2 predicts forms a β-hairpin (Fig. 1B, C) (Jumper *et al*, 2021). The TM2-TM3 loop is only present in organellar isoforms NHE6, NHE7 and NHE9 (Fig. 1C) and, similar to NhaA, multimer AlphaFold2 predicts the β-hairpins will interact with each other at the dimerization interface (Fig. 1D) (Jumper *et al*., 2021), but this region was too dynamic to be modelled in previous cryo EM structures (Fig. 1E) (Winklemann *et al*., 2020). Although, it is currently unclear if the TM2-TM3 loop domain contributes to lipid binding, the mutation of two lysine residues Lys105 and Lys107 in the unresolved TM2-TM3 loop domain, was sufficient to abolish PIP_2_ stabilization, indicating this might be the case (Winklemann *et al*., 2020).

**Figure 1.**
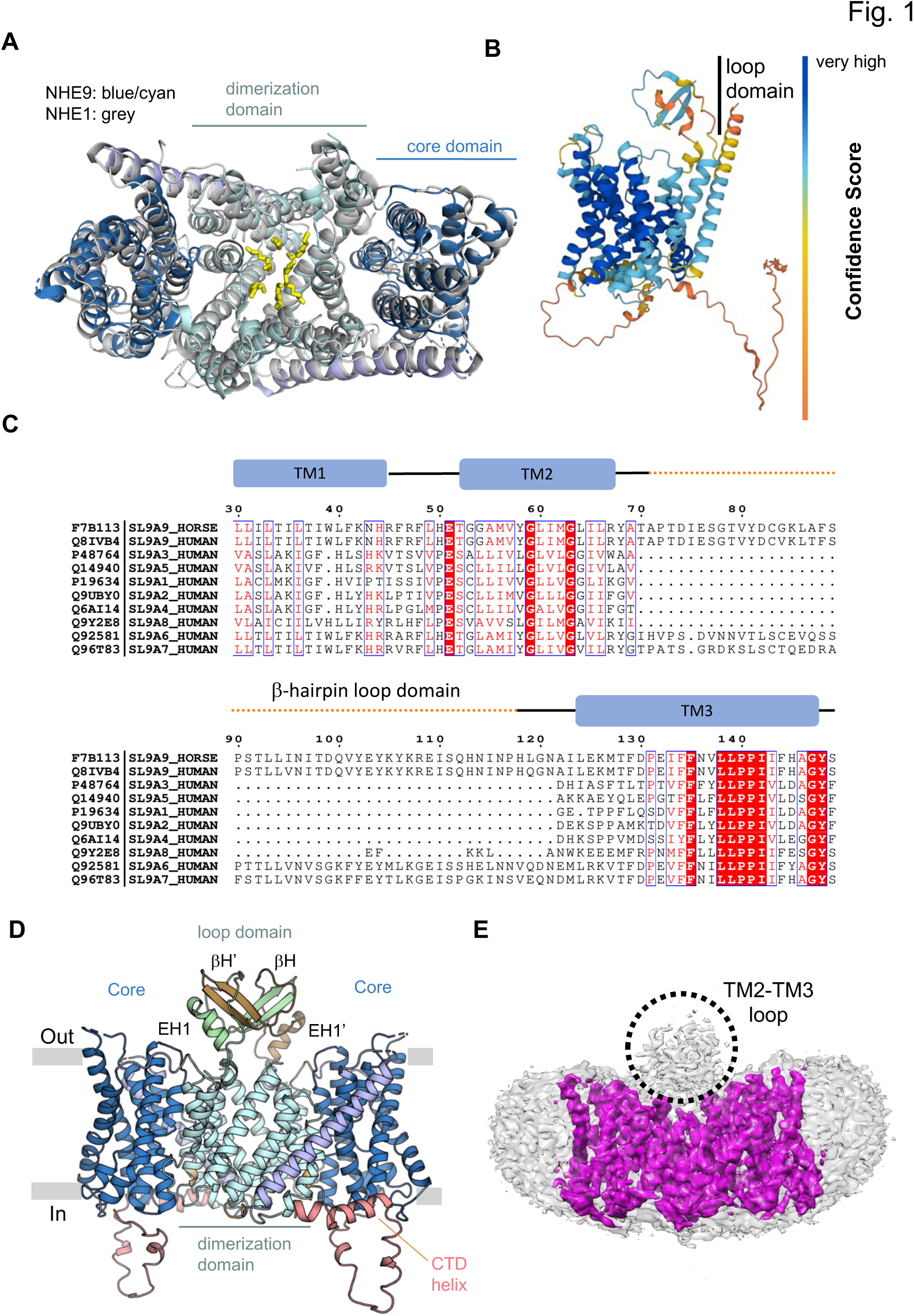
AlphaFold2 model for endosomal NHE9 includes β-hairpin TM2-TM3 loop domain. (**A**) Structural superimposition of *horse* NHE9* with inward-facing NHE1 (C_α_ RMSD = 2.91 Å) shows the overall structural conservation. Inward-facing NHE9 ΔCTD (PDB: 6Z3Z) with dimerization domain (cyan), core domain (blue) and TM7 linking helix (light-purple) all shown as cartoon. Inward-facing NHE1 (PDB: 7DSW) (grey, cartoon) with its lipids (yellow, sticks) inside the cavity between the dimerization domains. (**B**) AF2 model of the *human* NHE9 monomer that predicts an additional TM2-TM3 loop domain containing a ϕ3-hairpin with reasonable confidence. **(C)** The *human* NHE1-9 sequence alignment for TM1 to TM3. Residues with over 90% sequence identity are shown in red. The ϕ3-hairpin TM2-TM3 loop is only present in organellular NHE6, NHE7 and NHE9 proteins. (**D**). AF2 model of the *horse* NHE9 dimer with an TM2-TM3 loop domain containing an ϕ3-hairpin (ϕ3H1, ϕ3H1’) and extracellular helix (EH1, EH1’) from each of the two protomers coloured in brown and green, respectively. The core domain (dark-blue), dimerization domain (light-blue) and CTD domain (salmon) is further highlighted. (**E**) NHE9* structure and maps (PDB: 6Z3Z, EMD: EMD-11067) at low contour levels to highlight the loop domain map features that could not be modelled (black-dotted circle).

Here, we set out to determine cryo-EM structures of the β-hairpin containing NhaA and NHE9 Na^+^/H^+^ exchangers to resolve the proposed lipid-mediated oligomerization and to establish how lipid-binding can allosterically regulate exchange activity.

## Results

### Cryo EM structure of the NhaA homodimer at pH 7.5 bound to CDL

*E. coli* NhaA has become a model system for Na^+^/H^+^ exchangers (Padan, 2008; Winkelmann *et al*., 2022). Despite extensive biochemical, biophysical, and computational analysis, most structures of NhaA were crystallized as monomers (Hunte *et al*., 2005a; Winkelmann *et al*., 2022). The NhaA homodimer was been crystallized, but at 3.5 Å resolution it was unclear if lipids mediated dimerization (Lee *et al*, 2014). As such, a *Ec*NhaA WT-like triple mutant (A109T, Q277G, L296M) was purified in detergent supplemented with cardiolipin (CDL) for cryo EM studies. In brief, from optimized grids, a data-set of 14,329 movies was collected and, after processing, a cryo EM map at 3.3 Å resolution could be obtained, according to the gold-standard Fourier shell correlation (FSC) 0.143 criterion (Supplementary Fig. 1A-C, Table 1).

Similar to previous NhaA crystal structures, the cryo EM structure of NhaA adopts the inward-facing conformation (Fig. 2A) (Hunte *et al*, 2005b; Winkelmann *et al*., 2022). Between inactive pH 4 and partially-active pH 6.5, a conformational switch termed a pH sensor/gate allows hydrated Na^+^ ions to enter the cavity and controls the pH at which NhaA becomes activated (Fig. 2A) (Padan *et al*, 2004; Winkelmann *et al*., 2022). As such, the mutation of charged residues on the cytoplasmic surface can alter the pH of NhaA activation and mutation of surface histidine residues to alanine abolished activity altogether (Padan *et al*., 2004; Winkelmann *et al*., 2022). Similar to the monomeric crystal structure at pH 6.5 (Winkelmann *et al*., 2022), an intracellular cavity in the cryo EM structure at pH 7.5 has opened to enable Na^+^ accessibility to the ion-binding (Fig. 2A).

**Figure 2.**
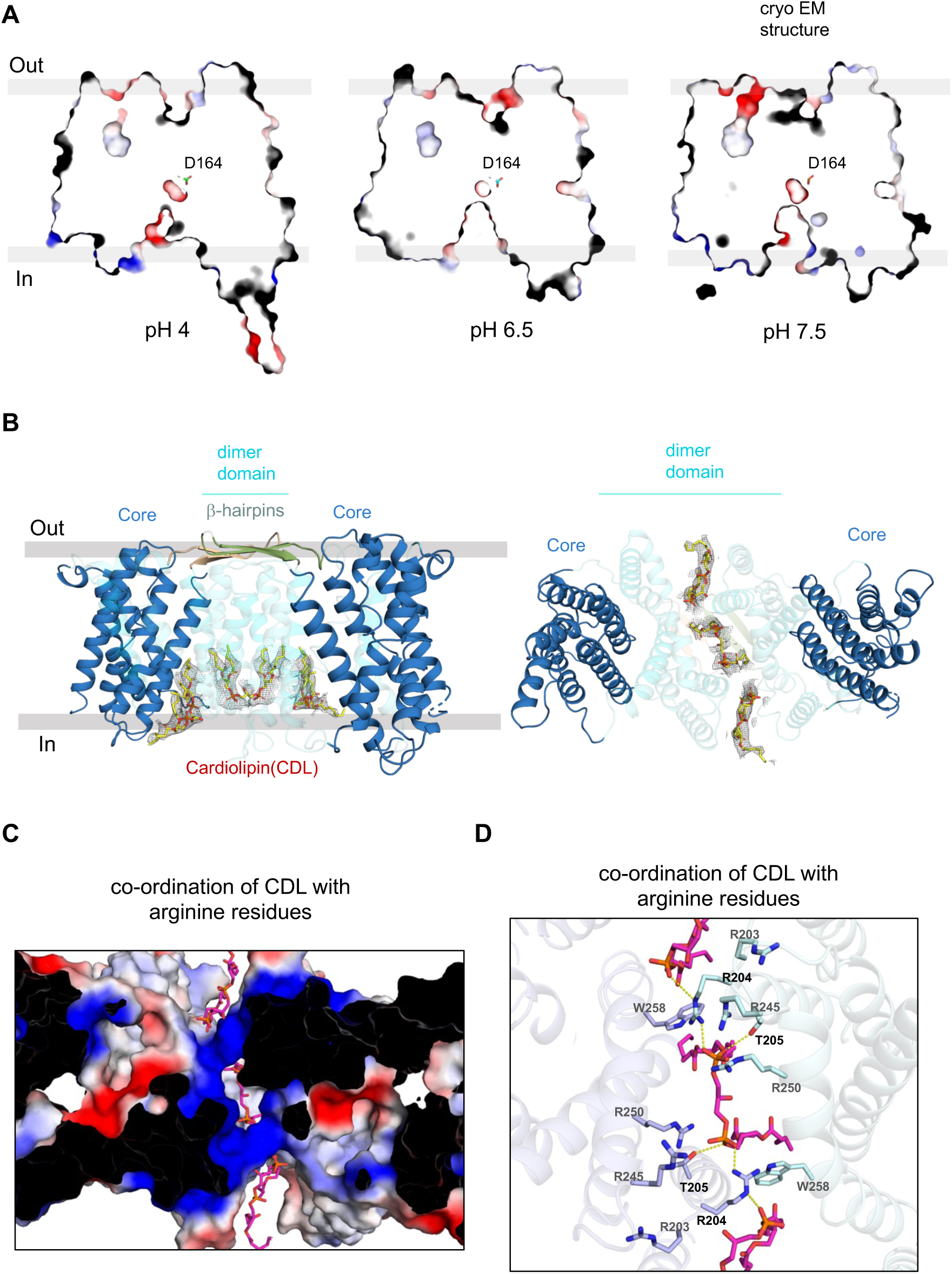
Structure of cardiolipin (CDL) bound to *Ec*NhaA homodimer. (**A**) Slice through an electrostatic surface representation of *Ec*NhaA structures, perpendicular to the membrane plane, at active pH 6.5 (left, PDB ID: 7S24), inactive pH 4.0 (right, PDB ID: 4AU5) and pH 7.5-CDL bound. The ion-binding aspartate is indicated and shown as cyan sticks. The ion-binding funnel at the cytoplasmic side is much more open at active pH 6.5 and pH 7.5-CDL bound structure than at pH 4. (**B**) *left:* Model of *Ec*NhaA homodimer obtained by cryo-EM and with β-hairpin loop domains (palegreen, sand, cartoon). Large densities between the dimerization domains indicate the presence of three CDL (yellow, sticks) fitted in the model. *right* Cytoplasmic view of *Ec*NhaA model with the three CDL fitted between the dimer domains. (**C**) Cytoplasmic view of slice through an electrostatic surface potential map with three cardiolipins (mangenta, sticks) positioned in the hydrophobic and positively-charged dimerization interface (blue). (**D**) Cytoplasmic view of *Ec*NhaA model (cartoon) with CDL (magenta, sticks) coordinating residues. Positively charged arginine residues are labelled and shown in cyan and purple sticks.

The cryo EM structure of NhaA retained the physiological homodimer (Fig. 2B). As expected from native MS and MD simulations (Gupta *et al*., 2017), thermal-shift assays (Nji *et al*., 2018), and functional activity analysis (Quick *et al*, 2021; Rimon *et al*., 2019) clear map density to supports the modelling of cardiolipin at the dimerization interface (Fig. 2B). One CDL was present in the middle of the positively-charged dimerization interface, flanked by two additional CDL lipids bound towards the ends of the dimer interface (Fig. 2C). Although this part of the dimerization interface is very positively-charged with four arginine side chains per protomer contributing — Arg203, Arg204, Arg245 and Arg250 — only the Arg204 residue directly interacts with the phosphate headgroups of the central CDL. Additional hydrogen bonds are contributed by Thr205 and Trp258 to oxygen atoms of CDL phosphoester bonds (Fig. 2D). Deeper in the cleft, the side chain of Trp258 hydrogen bonds to Arg204, and also interacts with both the central and the flanking CDL molecules, sandwiched between the distal glycerol moieties of both the middle and the flanking CDL lipids. The flanking CDL molecules hydrogen bonds to Arg204, and the backbone oxygens of Gly201, Ala202 and Arg203.

The co-ordination of CDL seen in *Ec*NhaA triple mutant is entirely consistent with computational analysis of CDL binding sites from analysis of more than 40 different *E. coli* proteins (Corey *et al*, 2021). Moreover, an Arg203Ala and Arg204Ala double-mutant abolished thermostabilization of NhaA by CDL (Winkelmann *et al*., 2022). The ion-binding site is located between cross-over helices in the core domain, yet accessibility to the ion-binding site is likely to be influenced by hydrophobic gating residues on the dimerization helix of TM2 (Matsuoka *et al*., 2022; Okazaki *et al*, 2019). It seems that stabilization of the scaffold by CDL might favour Na^+^ accessibility to the ion-binding site, which would be consistent with the higher affinity for ^22^Na^+^ as measured upon the addition of CDL to detergent-solubilised NhaA (Quick *et al*., 2021).

*Cryo-EM structure of NHE9* with TM2-TM3 β-hairpin loop domain at low pH 6.5* NHE9 was predicted by AlphaFold2 (Jumper *et al*, 2021) to harbour a β-hairpin loop domain between the topologically equivalent helices, TM2-TM3, to the β-hairpins in NhaA (Fig. 1D, Supplementary Fig. 2A). In the AlphaFold2 NHE9 monomer vs multimer prediction, the β-hairpins in NHE9 re-adjust their position to interact with the β-hairpin from the neighbouring protomer (Supplementary Fig. 2B). The previous structure of *horse* NHE9* (residues 8 to 574; out of 645) by cryo-EM was determined from protein purified in the detergents LMNG and CHS at pH 7.5, but map density was too poor to model this loop domain (Fig. 1E) (Winklemann *et al*., 2020). Here, we repeated the collection of NHE9* instead at pH 6.5 where NHE9 is thought to be less active, and therefore presumably less dynamic. Overall, we collected ∼4,300 movies and the final 3D-reconstruction contained data of ∼244,279 particles, from which an EM map could be reconstructed to 3.3 Å according to the gold-standard Fourier shell correlation (FSC) 0.143 criterion (Supplementary Fig. 3A). Comparison of the NHE9* structures determined at pH 7.5 and pH 6.5 showed no apparent differences (Supplementary Fig. 3B). We speculated that the respective final 3D reconstructions may still contain heterogeneity and we therefore performed 3D-variability analysis in CryoSPARC (Punjani *et al*, 2017), followed by clustering (principal component) analysis to separate further classes. Selecting 42% of the particles for heterogeneous refinement, enabled an improved cryo EM map reconstruction of NHE9* at pH 6.5, with additional map features for the TM2-TM3 loop domain (Supplementary Fig. 4). Further homogenous refinement with C2 symmetry improved the cryo EM density for the TM2-TM3 loop domain with an overall resolution estimate of 3.6 Å (Supplementary Fig. 4A), although the map resolution seems poorer at ∼ 4.2 Å resolution. As a last-step, a composite map was generated for model building, combining the additional TM2-TM3 loop domain features with the higher resolution EM maps (Supplementary Fig. 4B). Subsequently, we refined the AlphaFold2 NHE9 multimer model into the NHE9* cryo-EM maps with some further manual adjustment where needed (Methods and Fig. 3A-B).

**Figure 3.**
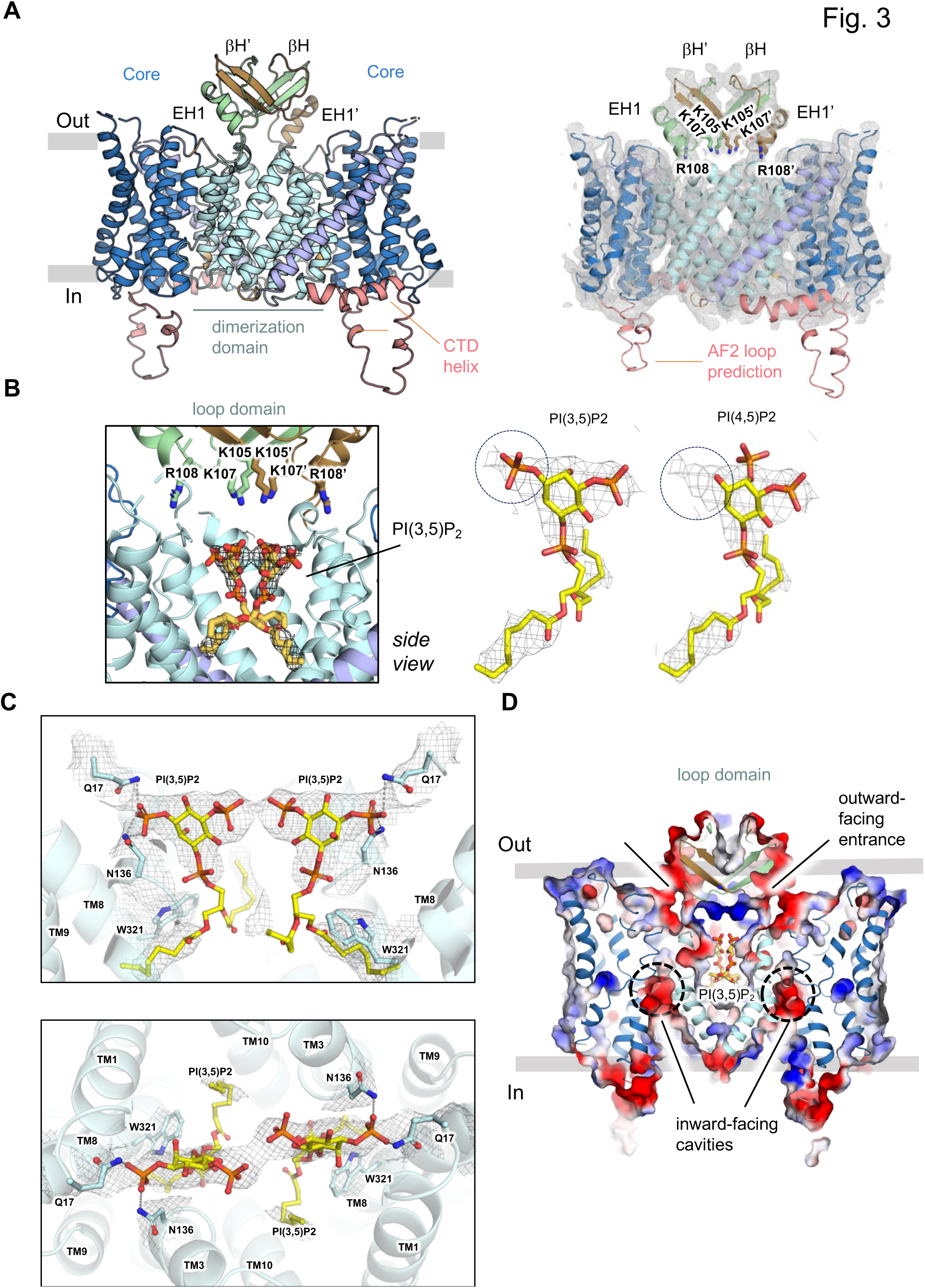
TM2-TM3 β-loop domain structure of NHE9* CC with negatively-charged PI(3,5)P_2_ lipids bound at the dimer interface. (**A**) *left:* Structure of the *horse* NHE9* structure determined by cryo-EM and guided by the refined NHE9* model for the β-hairpin TM2-TM3 loop domain. Domain-swapped βH, βH’ and EH1, EH1’ in green and brown respectively, and NHE9 core and dimer domains coloured as in Fig. 1D. An additional C-terminal helix in the CTD could be modelled (salmon, cartoon), but there is no density to support the 20 residue AF2 loop model that is located between the end of the core domain and the beginning of the interfacial helix *right:* The cryo EM maps for the *horse* NHE9* structure including the charged residues in loop domain highlighted in sticks. **(B)** *left:* In the *horse* NHE9* CC structure additional map density supported the modelling of two PIP_2_ lipids (yellow sticks) located in the middle at the dimerization interface between the two protomers and could interact with the positively charged β-hairpin TM2-TM3 loop domain residues (green and brown sticks). *right:* The fitting of PI(3,5)P_2_ vs PI(4,5)P_2_ lipids into the cryo EM maps are compared. (**C**) In addition to the β-hairpin TM2-TM3 loop domain lysine residues, PI(3,5)P2 is coordinated at the dimerization interface by aromatic and polar residues Trp321, Gln17, Asn136. TM segments TM1 and TM8 are not shown for better visualization of the bound lipids (yellow sticks and grey mesh) **(D)** Electrostatic surface potential map cross-section with two negatively-charged PI(3,5)P_2_ modelled in the hydrophobic and positively-charged dimerization interface (blue). The interface between the loop domains and core domains (indicated by the black-line) are negatively-charged and provide an electrostatic pathway for cations in the outward-facing state. Protein shown as cartoon colored as in Fig. 1E, and PI(3,5)P_2_ as yellow-sticks and the ion-binding site is highlighted (dotted-circle).

At the extracellular end of TM2, a flexible loop of residues Pro72 to Asp82, were predicted by AlphaFold2 as extending towards the centre of the dimerization interface, followed by short-linker and a β-hairpin strand angled 60° away from the NHE9* surface (Fig. 3A). We were able to confidently refine the overall position of the short linker and β-hairpin strand that fits the AlphaFold2 model (Fig. 3A). The loop domain reconnects to the beginning of TM3 *via* a short extracellular helix (ECH1), which was well supported by the cryo EM maps (Fig. 3A). The β-hairpins are predicted by AF2 to be domain-swapped, and this arrangement refines well into the map density. The centre of the loop domain has a cluster of positively-charged residues Lys105, Lys107 and Arg108 from each of the two protomers (Fig. 3A). Overall, the TM2-TM3 β-loop domain clasps the two protomers together on the luminal side, and forms a highly positively-charged cluster that is now located above the dimerization domain interface. As such, the re-modelled NHE9* structure is consistent with its previously proposed requirement for binding negatively-charged PIP_2_ lipids at the interface (Winklemann *et al*., 2020).

*Cryo-EM structure of cysteine variant (NHE9*CC) shows binding of PI(3,5)P*_2_ *lipids* To improve the resolution of the NHE9* structure we substituted Leu139 and Ile444 residues to cysteines in an attempt to disulphide-trap the inward-facing state. In addition, we added brain lipids during each step of the NHE9*CC affinity purification (see Methods). We subsequently collected a larger data-set of 13,780 movies and the final 3D-reconstruction had 78,370 particles from which an EM map was reconstructed to 3.2 Å according to the gold-standard Fourier shell correlation (FSC) 0.143 criterion (Supplementary Fig. 5A). Despite the similar FSC resolution estimate, these cryo-EM maps were improved with some local regions extending to 2.5 Å resolution (Supplementary Fig. 5A). In particular, the density was clearer for the TM2-TM3 β-loop at lower contour levels (Supplementary Fig. 5A). Map density and sulphur atom distances, however, did not support disulphide bond formation between the two introduced cysteine residues (Supplementary Fig. 6A-B). Currently, it is unclear whether the improved map quality for NHE9*CC is due to the introduced cysteine mutations, or by the repeated addition of lipids during purification.

At the dimerization interface we further observe additional lipid-like features in the improved cryo EM maps (Fig. 3B and Supplementary Fig. 6C). Rather than the cylinder-like density of fatty acids seen in the NHE1 nanodisc structure in POPC lipids (Dong *et al*, 2021), the cryo-EM maps show larger, head-group lipid densities with distinct features (Fig. 3B, Supplementary Fig. 6C). Given that NHE9* co-purified from yeast with several lipids corresponding to the molecular mass of PIP_2_ (Winklemann *et al*., 2020), we attempted to model two PIP_2_ molecules into the NHE9* CC maps and found that the density was a better fit for the lipid phosphatidylinositol-3,5-bisphosphate PI(3,5)P_2,_, rather than predominantly found phosphatidylinositol-4,5-bisphosphate PI(4,5)P_2_ lipids (Fig. 3B, Supplementary Fig. 6C) — although PI(3,5)P_2_ is a minor PIP_2_ lipid, it is specific to late endosomes and lysosomes, which overlaps with the functional localization of NHE9 (Hasegawa *et al*, 2017) whereas PI(4,5)P_2_ is principally found at the plasma membrane (Ho *et al*, 2012). The glycerol backbone and acyl chains of the PI(3,5)P_2_ lipid form hydrophobic stacking interactions to Trp321 (TM8, TM8’) (Fig. 3C). Two glutamine residue Gln17 (TM1, TM1’) and Asn136 (TM3, TM3’) are well positioned to hydrogen-bond with the phosphomonoester in the C3-OH position (Fig. 3C). The position of these polar residues could partially explain the preference for PI(3,5)P_2_ lipids, since there are no polar residues close enough to interact with a PO_4_^2-^ at the C4-OH position.

Masked refinement of NHE9*CC was required to obtain high resolution maps to model PI(3,5)P_2_ lipids, but map features for the TM2-TM3 β-hairpin loop domain could not be retained, likely as its too dynamic (Supplementary Fig. 5A). Nevertheless, at lower contour levels, the map density for the β-hairpin loop domain matches the map density seen in the NHE9* maps at pH 6.5 (Supplementary Fig. 5A). With a rigid-body fit of the TM2-TM3 β-hairpin loop domain modelled into the NHE9*CC maps, the positively-charged loop domain residues Lys105, Lys105’ Lys107, Lys107’ could neutralise the negatively-charge headgroups of PI(3,5)P_2_ (Fig. 3D). Consistently, mutation of these lysine residues to glutamine was previously shown to abolish PIP_2_ – induced thermostabilization of NHE9 (Winklemann *et al*., 2020).

To evaluate if the lysine residues could interact with the modelled PIP_2_ lipids as expected, we carried out molecular dynamics (MD) simulations of the NHE9*CC structure and a *in silico* model for an NHE9* Lys85Gln-Lys105Gln-Lys107Gln variant (see Methods). In MD simulations, the PI(3,5)P_2_ lipid stayed within less than 3 Å from its initial position for nearly the entire simulation time (Fig. 4A,B). In contrast, a modelled PI(4,5)P_2_ lipid was less stably bound with a higher fraction of frames showing > 3 Å movement from its initial position (Fig. 4B). The distribution of salt-bridge interactions with Lys105 and Lys107 residues revealed that Lys105 residues were nearly always in contact with the PI(3,5)P_2_ lipid, whereas Lys107 residues remained in contact for around half of the simulation time (Fig. 4C,E). The PI(4,5)P_2_ lipid had a much larger variation for interaction with the lysine residues and the preferred interaction with Lys105 was less pronounced (Fig. 4D, E). In MD simulations of the modelled NHE9* Lys85Gln-Lys105Gln-Lys107Gln variant, neither of the PIP_2_ lipids were stably bound and the lipids had no clear interaction with the modelled glutamines (Fig. 4B-D). Overall, the MD simulations support that lysine residues in the TM2-TM3 β-hairpin loop can form stable interactions with the modelled PI(3,5)P_2_ lipids observed in the NHE9*CC variant. Thus, the polar and aromatic residues located at the dimerization interface and together with positive-charged residues from the TM2-TM3 β-hairpin loop, create an environment well-suited for binding the endosomal specific PI(3,5)P_2_ lipid (Fig. 3D).

**Figure 4.**
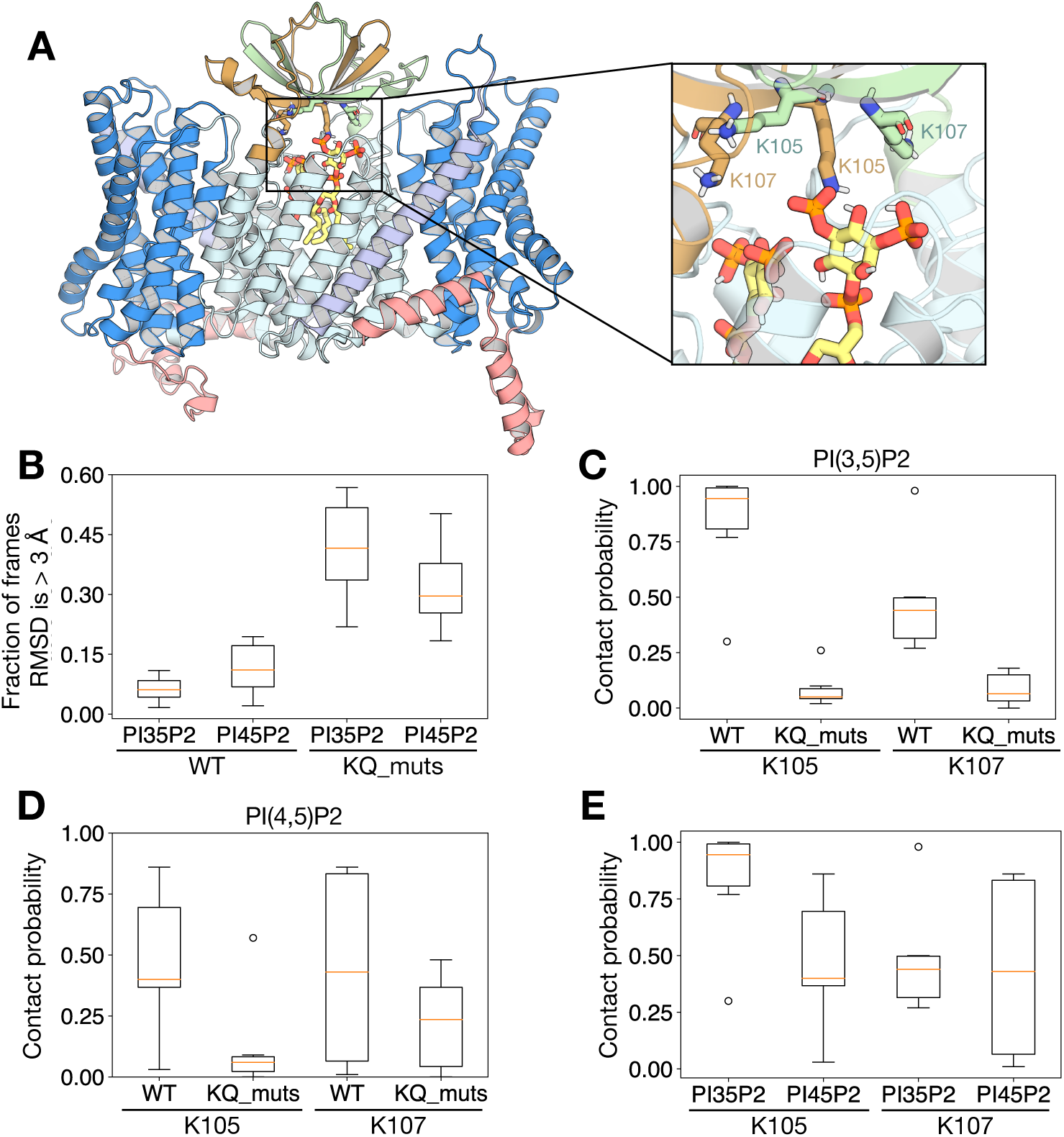
MD simulations of PIP_2_ interaction to NHE9* CC. (**A**) A representative snapshot of NHE9 with PI(3,5)P_2_ after a 250 ns simulation. The structure of PI(3,5)P_2_ is shown in yellow. K105 and K107 of the different protomers are shown in green and brown, respectively. Protein shown as cartoon and lipids as sticks (colored as in Fig. 1D). (**B**) Box plot representation of the distribution of the fractions of frames where the root mean square deviation (RMSD) of PIP_2_ lipids is > 3 Å with respect to its previous 10 ns structure. (**C**) Box plot representation of the distribution of frames where PI(3,5)P_2_ are within 4 Å (defined as contact probability) with either K105, K107 or its mutants. (**D**) Box plot representation of the distribution of frames where PI(4,5)P_2_ are within 4 Å (defined as contact probability) with either K105, K107 or its mutants. (**E**) Box plot representation of the distribution of the contact probability between K105 and K107 with either PI(3,5)P_2_ or PI(4,5)P_2_.

### Assessing the requirement of PI(3,5)P_2_ lipid binding to NHE9

We had previously probed interactions with PIP_2_ and PIP_3_ lipids to NHE9* using FSEC-TS and GFP-based thermal stability assays (Hattori *et al*, 2012; Nji *et al*., 2018; Winklemann *et al*., 2020), which we had earlier confirmed could detect cardiolipin-specific stabilization of *E. coli* NhaA (Landreh *et al*, 2017; Nji *et al*., 2018). The average melting temperature (Δ*T*_m_) of NHE9* increased by 8°C with PI(4,5)P_2_ addition, whereas other lipids POPC, POPE and POPA showed no clear thermostabilization (Winklemann *et al*., 2020). To validate that PI(3,5)P_2_ would also stabilise NHE9* in a similar manner, NHE9* GFP-TS melting curves were recorded in the presence of either PI(4,5)P_2_ or PI(3,5)P_2_ lipids (Methods and Fig. 5A). Consistently, NHE9* displayed higher thermostabilization with PI(3,5)P_2_ addition with a Δ*T*_m_ of 15°C, as compared to Δ*T*_m_ of 8°C for PI(4,5)P_2_ addition. Without lipid addition, NHE9* unfolds around 30°C with a shallow slope for the transition temperature, which is indicative of a mixed protein population (Fig. 5A). In contrast, with PI(3,5)P_2_ added, NHE9* melted with a sharper transition, indicating a shift to a more uniform protein species (Fig. 5A).

**Figure 5.**
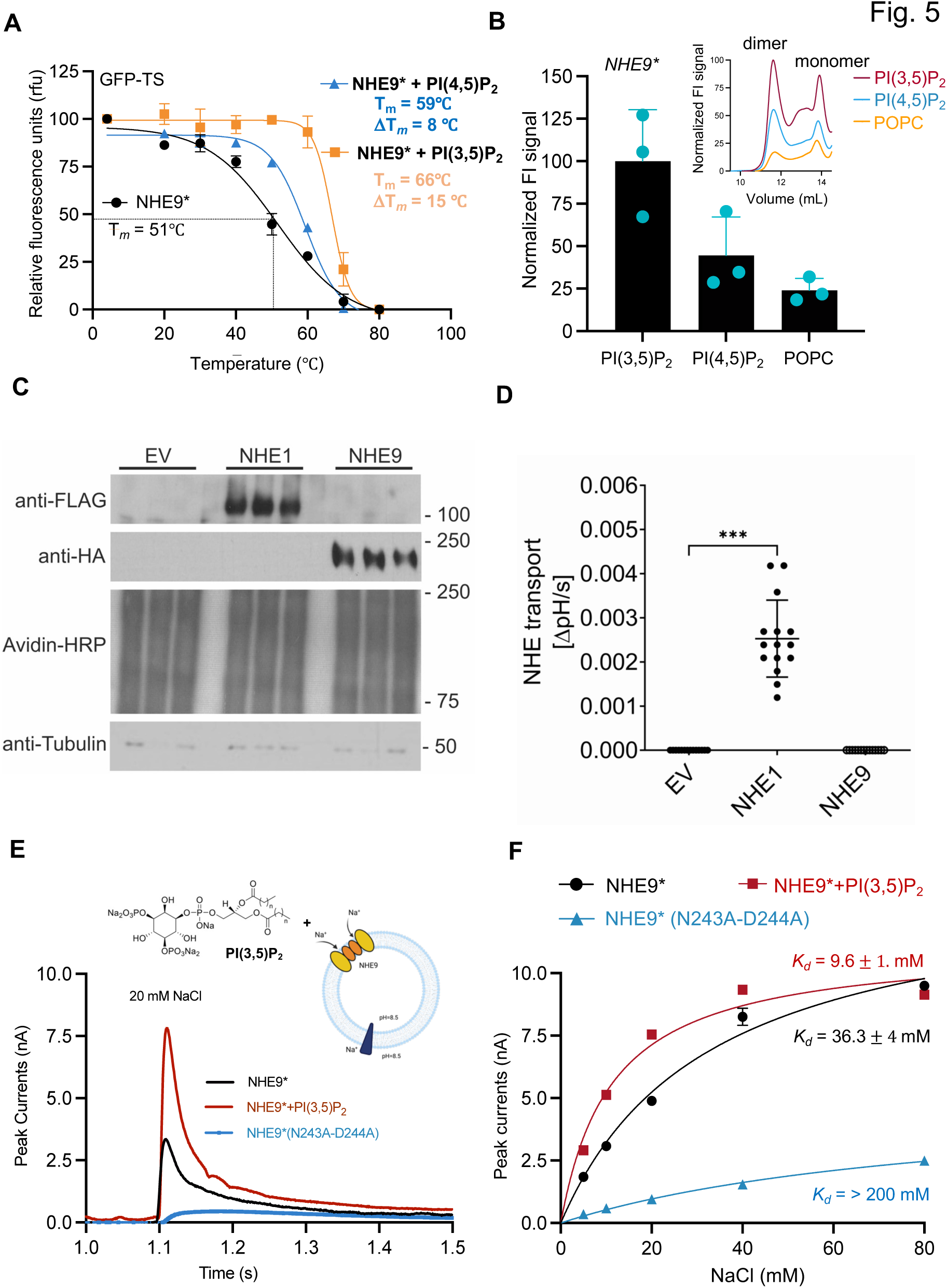
NHE9 is non-functional at the plasma membrane and the endosomal-specific lipid PI(3,5)P_2_ stabilizes the functional homodimer and activity. (A) Thermal shift stabilization of purified dimeric NHE9*-GFP in the presence of PI(4,5)P_2_ (blue) and PI(3,5)P_2_ (mustard) compared to lipid-free (black). Data presented are normalized fluorescence of mean values ± data range of n = 3 technical repeats; the apparent *T*_M_ was calculated from datapoints fitted according to a sigmoidal 4-parameter logistic regression function. (**B**) Bars represent the normalised fraction of NHE9* homodimer remaining after heating at 50°C for 10 mins in the presence of DDM-solublised lipids as labelled, which was asseded by FSEC. Error bars are the SEM of n = 3 FSEC injections, *top inset:* representative FSEC traces of the tabulated data shown. (**C**) Plasma membrane expression of FLAG-tagged NHE1 and HA-tagged NHE9 in PS120 cells. PS120 were transfected with empty vector (EV; pMH), human NHE1 or human NHE9. Plasma membrane proteins were isolated using surface biotinlyation. Avidin–HRP was used as loading control. The absence of a strong tubulin signal confirms the purity of the membrane fraction. Three independent biological replicates per condition. (**D**) NHE transport activity in transfected PS120 cells. Measurement of plasma membrane NHE transport activity in empty vector (EV; negative control), human NHE1 (positive control) or human NHE9 transfected PS120 cells by quantification of the sodium-dependent intracellular pH recovery after acidification of the cytoplasm. Values are shown as means ±SD. (**E**) SSM-based electrophysiology. Transient currents were recorded for NHE9* proteoliposomes (black trace) at symmetrical pH 7.5 after the addition of 20 mM NaCl. Peak currents for an ion-binding site NHE9* variant N243A-D244A (blue trace) and NHE9* pre-incubated with synthetic PI(3,5)P_2_ lipids (red trace) are also shown. *above:* schematic of NHE9* and PI(3,5)P_2_ reconstituted into liposomes for SSM based electrophysiology measurements (**F**) Fit of the amplitude of the transient currents as a function of Na^+^ concentrations at pH 7.5 for horse NHE9* proteoliposomes pre-incubated with either buffer (black-trace) or synthetic PI(3,5)P_2_ lipids (red trace) and the corresponding binding affinity (*K*_D_). Error bars are the mean values ± s.e.m. of: n = 3 individual repeats.

In the presence of various lipids, we further assessed the relationship between resistance to heat denaturation and retention of the NHE9 homodimer by FSEC (Methods). As anticipated, we observed that a higher fraction of the NHE9* homodimer was retained if PI(3,5)P_2_ lipid was added prior to heating (50°C for 10 mins), versus either PI(4,5)P_2_ or POPC lipid addition (Fig. 5B). We had previously shown that substitution of β-hairpin TM2-TM3 lysine residues to glutamine in NHE9* Lys85Gln-Lys105Gln-Lys107Gln abolished PI(4,5)P_2_ stabilization and oligomerization using native MS (Winklemann *et al*., 2020). Consistently, the NHE9* Lys85Gln-Lys105Gln-Lys107Gln variant purified as a monomer in detergent, and the addition of either PI(3,5)P_2_, PI(4,5)P_2_ or POPC lipids showed no clear thermostabilization (Supplementary Fig. 7A,B). Taken together, thermostability-shift assays confirms that NHE9* binds lipid PI(3,5)P_2_ and its addition stabilises the functional homodimer in detergent.

NHE9 is localized to endosomes (Nakamura *et al*, 2005). If PI(3,5)P2 lipids are required for NHE9 activity, then the protein might be inactive in membranes lacking this lipid. To assess the requirement of NHE9 activity for endosomal-specific lipids, we expressed *human* NHE9 and plasma membrane localized *human* NHE1 in a PS120 cell line, which is deficient in plasma membrane NHEs (Methods). In this mutant cell line, plasma membrane localization of both *human* NHE1 and NHE9 isoform was confirmed after their transient transfection by surface biotinylation, isolation and immunoprecipitation (Fig. 5C). To measure *human* NHE1 and NHE9 activity, the pH-sensitive dye BCECF was trapped intracellularly and pH recovery after acid loading with NH_4_Cl assessed spectroscopically. Whilst sodium-dependent intracellular pH recovery was clearly measurable in PS120 cells expressing *human* NHE1, no NHE activity could be measured for *human* NHE9, despite similar expression levels (Fig. 5C and D). Thus, we can confirm that *human* NHE9 is inactive when mistargeted to the plasma membrane.

As it is practically infeasible to incorporate PI(3,5)P_2_ lipids into the plasma membrane of cultured cells, we required another approach to assess whether NHE9 activity is also dependent on PIP_2_ lipids. Previously, NHE9* was reconstituted into liposomes together with F_0_F_1_-ATP synthase for proteoliposome studies and an apparent *K*_M_ of NHE9* for Na^+^ of 20.5 ± 3 mM was determined (Winklemann *et al*., 2020). However, a high background from empty liposomes would make assessing lipid requirements for NHE9* challenging in this set-up, as the signal-to-noise ratio was previously low ∼ 3:1 and the bacterial ATPase protein has different lipid preferences to NHE9. More recently, we have used solid-supported membrane (SSM) electrophysiology to record Na^+^ translocation of SLC9B2 and SLC9C1 Na^+^/H^+^ exchangers (Matsuoka *et al*., 2022; Yeo *et al*., 2023). In this setup, proteoliposomes containing NHEs are adsorbed to a SSM and charge translocation is measured *via* capacitive coupling of the supporting membrane, following accumulation of transported ions (Bazzone *et al*, 2017).

We forthwith recorded SSM responses for NHE9* proteoliposomes upon the addition of 20 mM NaCl (Supplementary Fig. 7C and Methods). Peak currents were ∼6-fold higher than either the NHE9* variant for which ion-binding site residues Asn243 and Asp244 were substituted to alanine, or the unrelated mammalian transporter for fructose (GLUT5) (Supplementary Fig. 7C). The estimated binding affinity (K_d_) determined by SSM for Na^+^ for NHE9* was 36.3 ± 4 mM and NHE9*CC was 24.1 ± 8 mM, which were similar to the Michaelis–Menten *K*_M_ estimate for NHE9* in proteoliposomes (Supplementary Fig. 7D) (Winklemann *et al*., 2020). Thus, we confirm that we can monitor Na^+^ binding to NHE9* using SSM, and that the protein binds sodium ions at a similar concentration range to the *in vivo K*_M_ estimates measured for other intracellular localized NHE6 and NHE8 isoforms (Pedersen & Counillon, 2019).

To assess the influence of PI(3,5)P_2_ lipid addition to NHE9* activity, we incubated the purified NHE9* protein with either buffer, or buffer containing solubilised PI(3,5)P_2_ lipids and reconstituted the respective mixtures into liposomes made from yeast-polar lipids (Methods). SSM-based electrophysiology of NHE9* incubated with buffer produced similar peak currents upon Na^+^ addition to those shown previously, whereas NHE9* protein incubated with PI(3,5)P_2_ lipid produced a stronger response to Na^+^ addition (Fig. 5E-F). Consistently, we observed a four-fold increase in its binding affinity for Na^+^ (*K*_d_ = 9.6 mM) (Fig. 5F). In contrast, PI(4,5)P_2_ lipid addition had a similar Na^+^ binding affinity as the buffer only samples (Supplementary Fig. 7E). To confirm that the response to PI(3,5)P_2_ addition was mediated by the expected interaction with the lysine residues in the TM2-TM3 β-hairpin loop domain, we recorded SSM-based currents for the purified lysine-to-glutamine NHE9* Lys85Gln-Lys105Gln-Lys107Gln variant. The triple glutamine variant also showed a weaker affinity for Na^+^ (*K*_d_ = 55 mM) compared with NHE9* and, in fact, the addition of the PI(3,5)P_2_ lipid decreased its apparent affinity for Na^+^ (*K*_d_ = 88 mM) (Supplementary Fig. 7F). The outside surface of the PI(3,5)P_2_-stabilised NHE9* TM2-TM3 β-hairpin loop domain has a partially negatively-charged surface (Fig. 3D). Is seems that stabilization of the TM2-TM3 β-hairpin loop domain by PI(3,5)P_2_ lipids provides an electrostatic pathway that can better attract ions to the outward-facing funnel.

### Further mechanistic insights from the improved cryo-EM NHE9 CC* structure

In the improved NHE9*CC maps, we were able to confidently model all side-chains forming the ion-binding site, which are positioned around the half-helical cross-over (Fig. 6A and B). We observed some minor differences between the side-chain positions of Thr214, Asp215, Glu239 and Arg408 from the previously reported NHE9* structure (Winklemann *et al*., 2020), and also between NHE9 and NHE1 (Fig. 5B, C). In MD simulations of NHE9 it was observed that Na^+^ can be coordinated within the core domain forming interactions to Asp244, Asn243 Ser240 and Thr214 and several waters (Winklemann *et al*., 2020). Based on phylogenic analysis (Masrati *et al*, 2018), it was predicted that a salt-bridge would also form between Glu239 in TM6 and Arg408 in TM11. Here, we can confirm a salt-bridge is indeed formed between Glu239 and Arg408 residues and given the position of Glu239 in the ion-binding site, we propose this salt-bridge aids the stabilization of the residues required for coordinating Na^+^ (Fig. 6C). Moreover, Thr214 forms a parallel hydrogen bond to Asn243 (Fig. 6C). This unexpected hydrogen bond establishes a more rigid ion-binding site than that observed in NHE1, since the Thr214 residue is replaced by valine in NHE1 (Fig. 6B, C). Indeed, all the plasma membrane localized NHE isoforms have a hydrophobic residue in this position, whereas the intracellular isoforms have a threonine residue (Winklemann *et al*., 2020). It’s plausible that the structural differences might explain why intracellular isoforms are thought to be able to transport K^+^ in addition to Na^+^ (Donowitz *et al*, 2013; Pedersen & Counillon, 2019).

**Figure 6.**
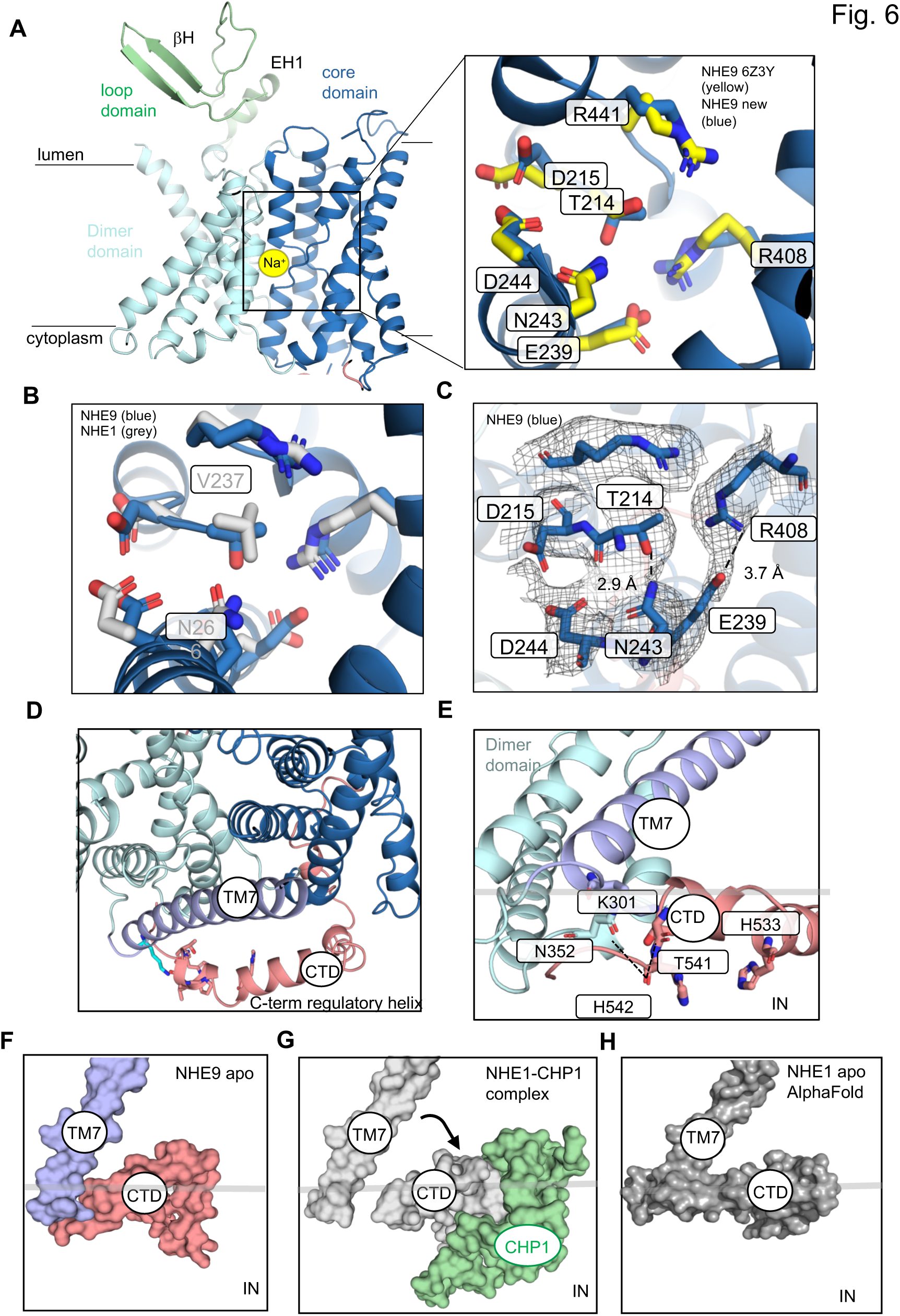
Ion-binding site and C-terminal domain of the NHE9*CC. (**A**) *left:* cartoon representation of the NHE9 ion-binding site, located in the 6-TM core transporter domain, which is made up of two broken helices. The sodium ion (yellow sphere) is located and coordinated at the ion-binding site. *right:* ion-binding residues of the NHE9*CC structure are shown as blue sticks and labelled, with the identical residues of NHE9* shown as yellow sticks (PDB id: 6Z3Y). **(B)** Ion-binding residues of NHE9*CC are shown as blue sticks and NHE1 residues shown as grey sticks with Val237 labelled**. (C)** Hydrogen-bonding between T214-N243 and salt-bridge interaction between E239-R408 in the NHE9*CC structure are illustrated by dashed lines and the cryo EM maps shown as grey mesh. (**D**) Top view showing how the CTD helix is positioned on the outside of the mobile TM7 linker domain. (**E**) Numerous contacts are formed between the CTD and the linker helix TM7 and in particular K301 (blue-sticks) in TM7 and the CTD helix (salmon sticks) and labelled. **(F)** In the horse NHE9* CC structure the CTD stays connected to the linker helix TM7 (protein shown as surface). **(G)** CHP1 (green, surface) binding to the CTD of NHE1 (grey, surface) moves it away from the linker helix TM7. (**H)** AF2 model of NHE1 predicts CTD interacts to TM7 as seen in the NHE9* CC cryo EM structure.

In NHE proteins, extrinsic factors bind to a large, intracellular C-terminal regulatory domain of ∼125 – 440 amino acids, which is only found in the mammalian proteins (Brett *et al*., 2005; Donowitz *et al*, 2009; Fuster & Alexander, 2014; Orlowski & Grinstein, 2004). In NHE1, removal of the regulatory domain results in a constitutionally active transporter (Wakabayashi *et al*, 1992). The C-terminal domain has been referred to as an allosteric regulatory subunit, which influences ion-exchange activity *via* interaction with many effectors (Lacroix *et al*, 2004) e.g., calmodulin (CaM) (Donowitz *et al*., 2013; Donowitz *et al*., 2009; Norholm *et al*, 2011; Odunewu-Aderibigbe & Fliegel, 2014; Slepkov *et al*, 2007). The recent human NHE1 structure in complex with Calcineurin B homologous protein 1 (CHP1) revealed an interaction with an interfacial α-helical stretch formed by residues 517 to 539 (Dong *et al*., 2021). However, CHP1 interacting with the interfacial helix was found to have no direct contacts with the transporter module itself, and is currently unclear how CHP1 binding increases NHE1 activity (Dong *et al*., 2021). Key to developing an allosteric model for extrinsic regulation is to obtain an NHE structure without complex partners. In the improved NHE9*CC structure, we could model part of the C-terminal domain without protein complex partners (Fig. 6D, Supplementary Fig. 6A). The interfacial helix in the CTD sits on the membrane interface and wraps around from the core domain to the linker helix TM7 (Fig. 6D). In particular, the highly-conserved Lys301 on the linker helix makes direct hydrogen bond interactions to carbonyl of Ile539, backbone amine of Thr541 and carbonyl of Thr541 (Fig. 6D-E and Supplementary Fig. 8). Given that the NHE1 and NHE9 structures superimpose well, apart from the position of the interfacial helix, it seems likely that the binding of CHP1 to the loop region, proceeding the interfacial helix, has driven its dissociation from the linker helix (Fig. 6F,G). Consistently, without CHP1 present, the AlphaFold2 model of NHE1 is similar to NHE9, with clear interactions observed between TM7 and the CTD helix (Fig. 6H). The cytoplasmic surface is also more positively charged with the CTD helix in the likely inhibited position, which could diminish the attraction of positively-charged ions to the inward-facing cavity (Supplementary Fig. 9A-C).

## Discussion

Elevator transporters are distinguished from rocker-switch and rocking bundle transporters by the fact that substrates are translocated by just one of the two domains (Drew & Boudker, 2016). The transport domain is able to move independently from the scaffold domain and carry the substrate across the membrane, as the scaffold domain does not participate in substrate binding. It seems in elevator proteins oligomerization is likely required so that the transporter domain can move effectively against the scaffold domain. In some elevator proteins, such as Na^+^-coupled glutamate transporters, the scaffold domain is extensive and, once formed, oligomerization is thought to be lipid insensitive. In the Na^+^/H^+^ exchanger family, however, the scaffold domain can be flexible and shows large structural variations (Winklemann *et al*., 2020), thus implying a role susceptible to regulation. Indeed, in the Na^+^/H^+^ exchanger NHA2 (SLC9B2) the additional N-terminal helix can readjust its position in the presence of yeast PI-lipids to make a more compact homodimer (Matsuoka *et al*., 2022). Moreover, monomeric mutants of NHA2 are inactive and indicate that lipid-remodelling of the homodimer is likely required for functional activity (Matsuoka *et al*., 2022), although this is yet to be firmly established.

Here, we have compared the lipid-mediated oligomerization in the Na^+^/H^+^ exchanger *E. coli* NhaA and the endosomal exchanger *horse* NHE9, which both harbour β-hairpins located between topologically equivalent helices in the scaffold domain. The scaffold β-hairpin loop domain is absent is bacterial Na^+^/H^+^ NapA (Lee *et al*., 2013) and NhaP (Paulino *et al*., 2014) members, and in mammalian NHEs localised to the plasma membrane (Dong *et al*., 2021; Dong *et al*, 2022). Moreover, the high structural-similarity between plasma membrane localised NHE1 and endosomal NHE9, implies that the additional TM2-TM3 β-hairpin loop domain has an additional, regulatory role. As expected, CDL binds between NhaA protomers and is stabilised by interaction with arginine residues. In addition, CDL coordination further requires a tryptophan residue located at the end of one of the scaffold helices. Consistently, in computational-based screening for CDL binding to bacterial proteins, a tryptophan residue seems to be a key requirement for ligand lipid-like interactions (Corey *et al*., 2021). In *horse* NHE9, we observe the PI(3,5)P2 lipid interaction also with a tryptophan residue to one of the scaffolding helices at the dimerization interface. Although positively-charged residues in the TM2-TM3 β-hairpin loop domain are required to stabilise the negatively-charged lipid, the coordination is further established by the tryptophan residue, as well as other polar residues at the dimer interface. In particular, in both structures the lipid interacts with residues in the topologically-equivalent scaffold helices, TM2 in NhaA and TM3 in NHE9, which further harbour highly-conserved hydrophobic gating residues (Winklemann *et al*., 2020). These hydrophobic gating residues are located opposite to the ion-binding site, and they are thought to prevent elevator transitions, unless the negatively-charged aspartate has bound its substrate ion (Winklemann *et al*., 2020). Thus, we propose a common allosteric coupling between ion-binding and lipid-stabilized oligomerization, whereby stabilization of the scaffold helix enhances Na^+^ binding. As yet, it is unclear whether catalytic turnover would also increase with scaffold stabilization.

The structure of NhaA in complex with CDL supports the proposed regulatory switch induced by salt-stress in *E. coli* (Nji *et al*., 2018). In NHE9, the specific binding of PI(3,5)P_2_ is consistent with its subcellular location, and the lipid could either enhance or completely turn on NHE9 activity by stabilizing the homodimer once it reaches late-endosomes (Fig. 7). Consistent with a regulatory switch, PI(3,5)P_2_ is a minor lipid that is not found in the plasma membrane, but only found in late endosomes and lysosomes (Hasegawa *et al*., 2017; Ho *et al*., 2012), which coincides with NHE9 localization. Moreover, the V-type ATPase is known to co-localise with NHE9, and work together to fine-tune organellular pH (Kondapalli *et al*., 2015). Consistent with the proposed model, the activity of the V-type ATPase is also increased in the presence of PI(3,5)P_2_ lipids (Li *et al*, 2014). Lastly, inhibition of enzymes required to produce PI(3,5)P_2_ lipids results in impaired epidermal growth factor receptor (EGFR) trafficking (de Lartigue *et al*, 2009). Indeed, NHE9 activity has been shown to be critical for EGFR sorting and turnover (Kondapalli *et al*., 2015). Since intracellular NHEs recycle through the plasma membrane they could, in principle, acidify the cell upon exposure to high Na^+^ levels, and yet vesicular acidification in NHE7-expressing cells has not been measured (Milosavljevic *et al*, 2014) and NHE9 was likewise found to be inactive at the plasma membrane. To the best of our knowledge, such a lipid-conditioned-activation model would be a novel regulatory mechanism for ion-transporters and SLCs in general.

**Figure 7.**
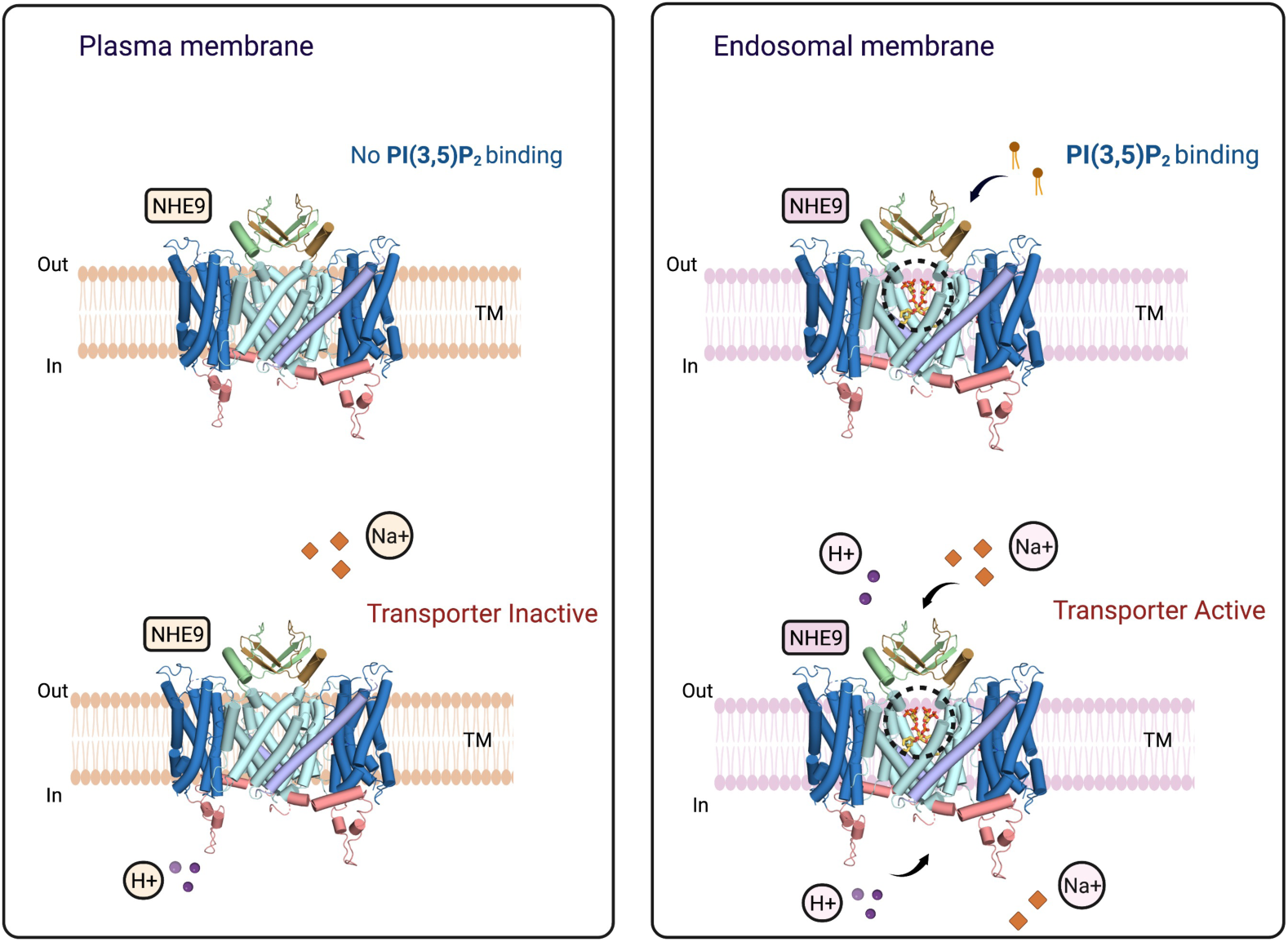
The PI(3,5)P_2_-dependent activation of NHE9 in late endosomes. Schematic representation of NHE9 ion translocation, dimer stabilisation and transport activation in the endosomes vs. the cell membrane. NHE9 is inactive in the plasma membrane. Upon relocation into late endosomes, a concomitant binding of PI(3,5)P_2_ at the dimerization interface improves the stability of homodimer and activates the transporter. The figure was created using Biorender.com

In addition to the lipid-dependent oligomerization, mammalian NHEs have a long C-terminal regulatory domain that regulates their activity (Pedersen & Counillon, 2019). Bacterial Na^+^/H^+^ exchangers, like NhaA, lack long C-terminal tails as presumably this level of extrinsic regulation is not required. The C-terminal domain is poorly conserved across the different NHE members and, so far, only part of the C-terminal tail could be modelled for *human* NHE1 in complex with CHP1 (Dong *et al*., 2021). Structures of human NHE1-CHP1 in outward and inward-facing states show some differences in the position of the C-terminal tail (Dong *et al*., 2021). Based on these structural differences, it was proposed that CHP1 may increase activity by favouring an outward-facing state. In the improved NHE9*CC structure we could model the corresponding region of the C-terminal tail, but in the absence of any interacting regulatory proteins. Surprisingly, we find that NHE9*CC has a similar interfacial helix between the core and dimer domains to that seen in NHE1. However, in the absence of a regulatory protein, the C-terminal interfacial helix in NHE9*CC is making a number of direct contacts with the linker helix TM7 through interaction with a lysine residue, likely restricting its mobility and that of the core transport domains. An autoinhibitory role of the C-terminal tail has been previously proposed for NHE1 (Wakabayashi *et al*, 1997), which is removed upon Ca^2+^-calmodulin binding to a site distal to the CTD helix (Sjogaard-Frich *et al*, 2021). Comparing our structure of NHE9 with NHE1-CHP provides a mechanistic framework for the autoinhibitory model, in that the relocation of an auto-inhibitory C-terminal tail by binding partners would remove the brake and enable ion-exchange.

## Summary

Our work provides a structural and molecular framework for the allosteric regulation of NhaA and NHE9 Na^+^/H^+^ exchangers by modulating oligomerization in a specific, lipid-dependent manner. Furthermore, we outline an activation model in the NHEs based on detachment of an CTD helix by accessory factors. Recently, it was shown that plant hormone (auxin) PIN-formed transporters are functional homodimers sharing the same fold as the Na^+^/H^+^ exchangers (Su *et al*, 2022; Ung *et al*, 2022; Yang *et al*, 2022). PIN transporters likewise harbour a C-terminal regulatory domain of varying length (Krecek *et al*, 2009), and also have both organellar and plasma membrane isoforms (Mravec *et al*, 2009), which must be regulated individually. Moreover, it has been shown that the plasma membrane localized PIN1 is in a dynamic equilibrium between monomers and dimers, and that the dimeric form can be regulated in the plant cell by endogenous flavonols (Teale *et al*, 2021). Thus, the allosteric regulatory mechanisms shown here by lipids and C-terminal tails, could reveal themes relevant to other transporters in general.

## Materials and Methods

### Expression and purification of NhaA and NHE9* and its variants

#### Expression and purification of EcNhaA-mut2

*Ec*NhaA WT-like triple mutant (A109T, Q277G, L296M), with a TEV-cleavable C-terminal GFP-His_8_ tag was overexpressed in the *E. coli* strain Lemo21 (DE3) and purified as previously described (Lee *et al*., 2014). Briefly, the *Ec*NhaA triple mutant was extracted from membranes with *n*-Dodecyl β-D-maltoside (DDM; Glycon) and purified by Ni-nitrilotriacetic acid (Ni-NTA; Qiagen) affinity chromatography. To purified NhaA-triple-mutant-GFP fusion, a final concentration of 3 mM cardiolipin (18:1) in 0.05% DDM was added, and then dialyzed overnight against buffer consisting of 20 mM Tris-HCl pH 7.5, 150 mM NaCl and 0.03% DDM. The dialysed *Ec*NhaA-triple-mutant-GFP fusion was subjected to size-exclusion chromatography and the peak concentrated to 3.5 mg.ml^-1^.

#### Expression and purification of horse NHE9*

The *horse* NHE9* structural construct (UniProt accession: F7B133) was identified previously (Winklemann *et al*., 2020), and is partially truncated on the C-terminal tail consisting of residues 8 to 575 out of a total of 644. The constructs NHE9*CC the L139C-I444C double mutant, NHE9*(N243A-D244A), NHE9*(K85Q-K105Q-K107Q) were synthesized and cloned into the GAL1 inducible TEV-site containing GFP-TwinStrep-His_8_ vector pDDGFP_3_. The cloned *horse* NHE9* and its variants were transformed into the *S. cerevisiae* strain FGY217 and cultivated in 24-L cultures of minus URA media with 0.2% of glucose at 30°C at 150 RPM Tuner shaker flasks using Innova 44R incubators (New Brunswick). Upon reaching an OD_600_ of 0.6 AU galactose was added to a final concentration of 2% (w/v) to induce protein overexpression. Following incubation at the same conditions the cells were harvested 22h after induction by centrifugation (5,000 ⨉ *g*, 4°C, 10 min). The cells were resuspended in cell resuspension buffer (CRB, 50 mM Tris-HCl pH 7.6, 1 mM EDTA, 0.6 M sorbitol) and subsequently lysed by mechanical disruption as previously described (Drew *et al*, 2008). Centrifugation (10,000 ⨉ *g*, 4°C, 10 min) was used to remove cell debris. Membranes were subsequently isolated from the supernatant by ultracentrifugation (195,000 ⨉ *g*, 4°C, 2 h), resuspended and homogenized in membrane resuspension buffer (MRB 20 mM Tris-HCl pH 7.5, 0.3 M sucrose, 0.1 mM CaCl_2_).

For structural studies of NHE9* at pH 6.5, the membranes were extracted and purified as described previously (Winklemann *et al*., 2020). In short, the Streptag-purified protein after removal of the C-terminal affinity-GFP-tag was collected and concentrated using 100 kDa MW cut-off spin concentrator (Amicon Merck-Millipore) further purified by size-exclusion chromatography (SEC), using a Superose 6 increase 10/300 column (GE Healthcare) and an Agilent LC-1220 system in 20 mM Mes-Tris pH 6.5, 150 mM NaCl, 0.003% (w/v) LMNG, 0.0006% (w/v) CHS. For NHE9*CC, the isolated membranes were solubilized as mentioned before (Winklemann *et al*., 2020) with addition of brain extract from bovine brain type VII (Sigma-Aldrich, cat. nr. B3635) to a total concentration of 0.003mg/ml, in the solublization, wash and elution buffers (Winklemann *et al*., 2020). The cleaved protein was collected and concentrated using 100 kDa MW cut-off spin concentrators (Amicon Merck-Millipore) separated by size-exclusion chromatography (SEC), using a Superose 6 increase 10/300 column (GE Healthcare) and an Agilent LC-1220 system in 20 mM Tris-HCl pH 7.5, 150 mM NaCl, 0.003% (w/v) LMNG, 0.0006% (w/v) CHS.

#### Cryo-EM sample preparation and data acquisition and processing

3 μg of purified NHE9*CC sample was applied to freshly glow-discharged Quantifoil R2/1 Cu300 mesh grids (Electron Microscopy Sciences) and blotted for 3.0s with a 20s waiting time prior, under 100% humidity and subsequently plunge frozen in liquid ethane using a Vitrobot Mark IV (Thermo Fisher Scientific). Cryo-EM datasets were collected on a Titan Krios G3i electron microscope operated at 300 kV equipped with a GIF (Gatan) and a K3 Bioquantum direct electron detector (Gatan) in counting mode. The movie stacks were collected at 130,000× magnification corresponding to a pixel size of 0.66 Å in a counted super-resolution mode. All movies were recorded with a defocus range of –0.4 to –2.5 µm. Similarly, 2.4 μg of purified NHE9* at pH 6.5 was blotted and the movie stacks were collected at 165,000x magnification, with a pixel size of 0.82 Å on Titan Krios G2 electron microscope operated at 300 kV equipped with a GIF (Gatan) and a K2 summit direct electron detector (Gatan) in counting mode The movies were recorded with a defocus range of –0.9 to –2.3 µm. 3µl of *Ec*NhaA-mut2 with concentration of 3.5mg/ml was blotted and the movie stack was collected at 130,000x magnification, with a pixel size of 0.6645 Å. The movies were recorded with a defocus range of –0.6 to –2.0 µm. The statistics of all cryo-EM data acquisition are summarized in Supplementary Table 1.

#### Image processing NHE9* at pH 6.5

The dose-fractioned movies were corrected by using MotionCorr2 (Zheng *et al*, 2017). The dose-weighted micrographs were used for contrast-transfer-function estimation by CTFFIND-4.1.13 (Rohou & Grigorieff, 2015). The dose-weighted images were used for auto-picking, classification and 3D reconstruction. Approximately 1,000 particles were manually picked, followed by a round of 2D classification to generate templates for a subsequent round of auto-picking using RELION-3.0 beta (Zivanov *et al*, 2018). The auto-picked particles were subjected to multiple rounds of 2D classification using RELION-3.0 beta to remove bad particles and “junk”. The particles belonging to “good” 2D classes were extracted and used for initial model generation using RELION-3.0 beta (Zivanov *et al*., 2018).

To visualize the extracellular loop domain of NHE9, the aligned 244,279 particles from RELION was imported into CryoSPARC (Punjani *et al*., 2017). *UCSF pyem (Daniel Asarnow, 2019)* was used for file format conversion from RELION to CryoSPARC. 3D Variability Analysis (3DVA) in CryoSPARC (Punjani *et al*., 2017) was performed and set up 6 variable components with the filter resolution 4 Å and high-pass filter 20 Å, respectively. After cluster analysis, the remaining 103,815 particles in 1 of 3 cluster was selected and subsequently, homogeneous refinement in CryoSPARC was performed after applying C2 symmetry and the reconstructed map reached to 3.1 Å resoltuion at the gold standard FSC (0.143).

#### Image processing NHE9*CC

The dataset was processed using CryoSPARC (Punjani *et al*., 2017). Dose fractionated movie frames were aligned using “patch motion correction,” and contrast transfer function (CTF) were estimated using “Patch CTF estimation”. The particles were picked using automated blob picker. Particles with good 2D classes were used for template based particle picking and extracted using a box size of 300 pixels. 268,054 particles were further used for *ab initio* model building and hetero refinements. Final round of non-uniform refinement with C2 symmetry and masked local refinement did result in 3D reconstruction with a gold standard FSC resolution estimation of 3.15Å.

#### Image processing EcNhaA-mut2

14,329 dose fractionated movie frames were aligned using “patch motion correction,” and contrast transfer function (CTF) were estimated using “Patch CTF estimation” in CryoSPARC (Punjani *et al*., 2017). 4,401,429 particles were extracted and cleaned up using multiple rounds of 2D classification. 542,404 particles were used for *ab initio* model building and cleaned using multiple rounds of hetero refinement. 78,917 particles were further selected for a final round of non-uniform refinement and masked local refinement which resulted in 3D reconstruction with a gold standard FSC resolution estimation of 3.37Å.

#### Cryo EM model building and refinement

Previously determined NHE9 structure (PDB id: 6Z3Z) was fitted into the cryoEM density map of NHE9* CC mutant. Iterative model building and real space refinement was performed using COOT (Emsley *et al*, 2010) and PHENIX.refine (Afonine *et al*, 2018). Prior to model building of the extracellular loop domain, AlphaFold2 (AF2) with poly-glycine linker was performed to predict the NHE9 dimer structure assembly. After automatically fitting the initial model into the cryo-EM map, iterative model building and real space refinement was performed using COOT (Emsley *et al*., 2010) and PHENIX.refine (Afonine *et al*., 2018). The refinement statistics are summarized in Supplementary Table 1. For generating structural figures PyMOL was used, and for figures of cryo-EM maps either Chimera (Pettersen *et al*, 2004) or ChimeraX (Pettersen *et al*, 2021) was used.

#### All-atom molecular dynamics simulations

The NHE9* CC structure and NHE9* TM2-TM3 loop variant (K58Q, K105Q, K107Q) were embedded into the POPC bilayer and solvated in 0.15 M NaCl using CHARMM-GUI (Qi *et al*, 2015). Either PI(3,5)P_2_ and PI(4,5)P_2_ were placed in the PIP_2_ binding site identified from the cryo-EM structure. All simulations were simulated with a 2 fs timestep using the CHARMM36m forcefield in GROMACS 2022.1(Abraham, 2015). The system was then energy minimised and equilibrated using the standard CHARMM-GUI protocol (Abraham, 2015), where the last step of the protocol was extended to 5 ns. The production runs were conducted for 250 ns under 303.15 K using the v-rescale thermostat. The pressure of all systems was maintained at 1 bar using a C-rescale barostat. All simulations were carried out in triplicates, where simulation frames were saved every 0.1 ns.

#### GFP-based thermal shift assay

To characterize lipid binding for NHE9* we used the GFP-Thermal Shift assay (Chatzikyriakidou *et al*, 2021). In brief, purified NHE9*-GFP fusions was isolated as described previously (Winklemann *et al*., 2020). A buffer containing 20mM Tris, 150mM NaCl and 0.03% (w/v) DDM 0.006% (w/v) CHS was used to dilute samples to a final concentration of 0.05–0.075 mg/ml with DDM added to a final concentration of 1% (w/v), and incubated for 30 min at 4°C. Stock solutions of the respective lipids, phosphatidylinositol-bis-4,5-phosphate (PI(4,5)P_2_ dioctanoyl Echelon Biosciences cat no. P-4508), and phosphatidylinositol-bis-3,5-phosphate (PI(3,5)P_2_ dipalmitoyl, Echelon Biosciences cat no. P-3516) were prepared with final concentration of 1 mg/ml in buffer mentioned above. Respective lipids were added to purified NHE9* sample to a final concentration of 0.1 mg/ml and incubated for 10 mins on ice. Subsequently, β-D-Octyl glucoside (Anatrace) was added to a final concentration of 1% (w/v) in 300 µl and the sample aliquots of 100 μl were heated at individual temperatures ranging from 20-80°C for 10 min using a PCR thermocycler (Veriti, Applied Biosystems). Heat-denatured material was pelleted at 5000 × g for 30 min at 4°C. The resulting supernatants were collected and fluorescence values recorded (Excitation: 488, Emission 512 nm) using 96-well plate (Thermo Fisher Scientific) measured with Fluoroskan microplate fluorometer (Thermo Scientific) reader. The apparent *T*m was calculated by plotting the average GFP fluorescence intensity from three technical repeats per temperature and fitting a resulting curve using a sigmoidal 4-parameter logistic regression using GraphPad Prism software (GraphPad Software Inc.). The Δ*T*_M_ was calculated by subtracting the apparent *T*_M_ of the lipid free sample, from the apparent *T*_M_ of the sample with the respective lipid added.

### Cell-based human NHE1 and NHE9 activity

#### Cell culture, constructs and transfection

Cells used for experiments were tested mycoplasma free and grown in a humidified 95/5% air/CO2-atmosphere incubator at 37 °C. PS120 fibroblasts, NHE1-null cells derived from Chinese hamster lung fibroblasts CCL39 cells (Pouyssegur *et al*, 1984), were cultured in high glucose DMEM supplemented with 10% fetal bovine serum (FBS), 100 units/mL penicillin and 100 µg/mL streptomycin (reagents from Sigma-Aldrich). Human full length NHE9 was cloned into the pMH vector (Roche) containing a C-terminal HA tag, human full length NHE1 was cloned into p3xFLAG-CMV-14 (Sigma-Aldrich) containing a C-terminal 3xFLAG tag. All constructs were verified by sequencing. Transient transfection was performed with Lipofectamine 2000 (Invitrogen) according to manufacturer’s instructions, and experiments (cell surface expression and transport function studies) were performed 48 hrs after transfection.

#### Cell surface biotinylation

Cell surface biotinylation was performed as described previously (Simonin & Fuster, 2010). Cells were rinsed with 1x PBS and surface proteins were biotinylated by incubating cells with 1.5 mg/ml sulfo-NHS-LC-biotin in 10 mm triethanolamine (pH 7.4), 1 mm MgCl_2_, 2 mm CaCl_2_, and 150 mm NaCl for 90 min with horizontal motion at 4 °C. After labeling, plates were washed with quenching buffer (1xPBS containing 1 mm MgCl_2_, 0.1 mm CaCl_2_, and 100 mm glycine) for 20 min at 4 °C, then rinsed once with 1xPBS. Cells were then lysed in RIPA buffer (150 mm NaCl, 50 mm Tris·HCl (pH 7.4), 5 mm EDTA, 1% Triton X-100, 0.5% deoxycholate, and 0.1% SDS), and lysates were cleared by centrifugation. Cell lysates of equivalent amounts of protein were equilibrated overnight with streptavidin-agarose beads at 4 °C. Beads were washed sequentially with solutions A (50 mm Tris·HCl (pH 7.4), 100 mm NaCl, and 5 mm EDTA) three times, B (50 mm Tris·HCl (pH 7.4) and 500 mm NaCl) two times, and C (50 mm Tris·HCl, pH 7.4) once. Biotinylated proteins were then released by heating to 95 °C with 2.5× Lämmli buffer.

#### SDS-PAGE, immunoblotting and antibodies

SDS-PAGE and immunoblotting were performed as described (Anderegg *et al*, 2020). Cell lysates were separated by SDS-PAGE and transferred to PVDF membranes. The membranes were incubated for 60 minutes with PBS-T20 containing 5% (w/v) skimmed milk. The membranes were then immunoblotted in 5% (w/v) skimmed milk in PBS-T20 with the indicated primary antibodies overnight at 4°C. The blots were then washed three to six times with PBS-T20 and incubated for 1 hour at room temperature with the appropriate secondary HRP-conjugated antibodies in 5% (w/v) skimmed milk in PBS-T20. After repeating the washing steps, the signal was detected with the enhanced chemiluminescence reagent. Immunoblots were developed using a film automatic processor (Fujifilm) and films were scanned with a 600-dpi resolution on a scanner and quantified with ImageJ software. Mouse monoclonal anti-FLAG M2 (#F3165), anti-HA (#H9658) and anti-Tubulin (#T9028) antibodies and were obtained from Sigma-Aldrich and used at 1:1000 dilution. HRP-coupled goat anti-mouse IgG antibody (#31430; Invitrogen) was used at 1:2000 dilution, avidin-HRP (#1706528; Bio-Rad) was used at 1:1000 dilution.

#### NHE1 and NHE9 transport activity

NHE activity was measured fluorometrically using the intracellularly trapped pH-sensitive dye BCECF (2ʹ,7ʹ-bis-(Carboxyethyl)-5(6ʹ)-carboxyfluorescein Acetoxymethyl Ester) (#B1170; Invitrogen) with the NH_4_Cl prepulse technique as described previously (Simonin & Fuster, 2010). Cells grown on glass coverslips were loaded with 1 μM BCECF-AM and exposed to 25 mM NH_4_Cl for 15 min in a buffer containing in mM: 120 NaCl, 5 KCl, 2 CaCl_2_, 1.5 MgCl_2_, 25 NH_4_Cl, 30 Hepes titrated to pH 7.4 with N-methyl-D-glucamine (NMDG). Then, cells were washed 3x with and incubated in a buffer containing in mM: 120 Tetramethylammonium-Cl (TMACl), 5 KCl, 2 CaCl_2_, 1.5 MgCl_2_, 30 Hepes titrated to pH 7.4 with NMDG. Recording was started and after 60 s cells were rapidly exposed to a buffer containing in mM: 120 NaCl, 5 KCl, 2 CaCl_2_, 1.5 MgCl_2_, 30 Hepes titrated to pH 7.4 with NMDG. BCECF fluorescence signals (λ excitation: 490 and 440 nm, λ emission: 535 nm) were recorded in a computer-controlled spectrofluorometer (Fluoromax-2, Photon Technology International). The 490/440 nm fluorescence ratio was calibrated to pHi using the K^+^/nigericin method, and initial rate (Vmax; βpH/βtime) of sodium-dependent intracellular pH recovery calculated. (Simonin & Fuster, 2010). All steps of incubation, recording and calibration were performed at 37 °C.

#### Solid Supported Membrane-based electrophysiology

For SSM-based electrophysiology measurements, protein was reconstituted in yeast polar lipids (190001C-100 mg Avanti). The lipids were prepared by solubilization in chloroform and dried using a rotary evaporator (Hei-Vap Core, Heidolph Instruments). Dry yeast polar lipids were thoroughly resuspended in 10 mM MES-Tris pH 8.5, 10 mM MgCl_2_ buffer at a final concentration of 10 mg ml^−1^. Unilamellar vesicles were prepared by extruding the resuspended lipids using an extruder (Avestin) with 200-nm polycarbonate filters (#10417004 Nuclepore Track-Etch membrane). The vesicles were destabilized by the addition of Na-cholate (0.65% w/v final concentration). SEC-purified protein was added to the destabilized liposomes at a lipid-to-protein ratio (LPR) of 5:1 and incubated for 5 min at room temperature. The sample was added to a PD SpinTrap G-25 desalting column (Cytiva) for removing detergent and the reconstituted proteoliposomes were collected in a final volume of 100μl. The sample was diluted to final lipid concentration of 5 mg ml^−1^ in 10mM MES-Tris pH 7.5, 10 mM MgCl_2_ buffer, flash frozen in liquid nitrogen and stored at −80 °C until use. Proteoliposomes were diluted 1:1 (vol/vol) with non-activating buffer (10 mM MES-Tris pH 7.5, 300 mM Choline chloride, 10 mM MgCl_2_) and sonicated using a bath sonicator. 10μl of sample was loaded on 1mm sensor (#161002 Nanion Technologies). For sample measured with PI(3,5)P_2,_ 20μM of the lipid final (v/v) was added to the protein and incubated for 15 min. The sample was subsequently reconstituted in yeast polar lipids as described above.

Sensor preparation for SSM-based electrophysiology using the SURFE^2^R N1(Nanion Technologies) system was performed as described previously (Bazzone *et al*., 2017). During the experiments, NHE9* and NHE9* mutants were activated by solution exchange from non-activating buffer to an activating buffer containing the substrate i.e NaCl. For binding kinetics, *x* mM Choline chloride was replaced by (300 – *x*) mM NaCl in the activating buffer at increasing concentrations. The Na^+^-induced peak currents were fitted from triplicate measurements using nonlinear regression curve-fit analysis to a one site specific binding model using GraphPad Prism software. The peak current values were normalized with respect to the average of the maximum value obtained across all measurements. The final *K*_d_ values reported are the mean ± s.d. of n = 3 independent sensors.

## Acknowledgements

We are grateful to Marta Carroni at the Cryo-EM Swedish National Facility at SciLife Stockholm for cryo-EM data collection as well as Michael Hall at the Umeå Core Facility for Electron Microscopy, UCEM and the European Synchrotron Radiation Facility (ESRF). The computations were enabled by resources provided by the Swedish National Infrastructure for Computing (SNIC) at the PDC Center for High Performance Computing, KTH Royal Institute of Technology, partially funded by the Swedish Research Council through grant agreement no. 2018-05973. This work was funded a European Research Council (ERC) Consolidator Grant EXCHANGE (Grant no. ERC-CoG-820187) to D.D

## Author contributions

D.D. designed the project. Cloning, expression screening and sample preparation for cryo-EM was carried out by P.M. and S.K. Cryo-EM data collection and map reconstruction was carried out by P.M, S.K, R.M, A.G an D.D. Model building was carried out by R.M, A.G and D.D. MD simulations were carried out by T.D and L.D. All authors discussed the results and commented on the manuscript. The authors declare no competing financial interests. Correspondence and request for materials should be addressed to D.D.

## Data availability

The coordinates and the maps for cryo-EM structures of *horse* NHE9* with TM2-TM3 loop at pH 6.5, *hor*se NHE9* CC bound to PI(3,5)P_2_, and the *E. coli* NhaA dimer with cardiolipin, have been deposited in the Protein Data Bank (PDB) and Electron Microscopy Data Bank (EMD) with entries PDB ID: 8PVR, 8PXB, 8PS0, respectively.

## References

1. The PyMOL Molecular Graphics System, Version 1.2r3pre ed. Schrödinger, LLC. Abraham MMT; Roland, S; Páll, S; Smith, J; Hess, B; Lindahl, E (2015) GROMACS: High performance molecular simulations through multi-level parallelism from laptops to supercomputers. SoftwareX 1: 19–25

2. Afonine PV, Poon BK, Read RJ, Sobolev OV, Terwilliger TC, Urzhumtsev A, Adams PD (2018) Real-space refinement in PHENIX for cryo-EM and crystallography. Acta Crystallogr D Struct Biol 74: 531–544

3. Anderegg MA, Albano G, Hanke D, Deisl C, Uehlinger DE, Brandt S, Bhardwaj R, Hediger MA, Fuster DG (2020) The sodium/proton exchanger NHA2 regulates blood pressure through a WNK4-NCC dependent pathway in the kidney. Kidney Int

4. Bazzone A, Barthmes M, Fendler K (2017) SSM-Based Electrophysiology for Transporter Research. Methods Enzymol 594: 31–83

5. Brett CL, Donowitz M, Rao R (2005) Evolutionary origins of eukaryotic sodium/proton exchangers. Am J Physiol Cell Physiol 288: C223–239

6. Chatzikyriakidou Y, Ahn DH, Nji E, Drew D (2021) The GFP thermal shift assay for screening ligand and lipid interactions to solute carrier transporters. Nat Protoc 16: 5357–5376

7. Coincon M, Uzdavinys P, Nji E, Dotson DL, Winkelmann I, Abdul-Hussein S, Cameron AD, Beckstein O, Drew D (2016) Crystal structures reveal the molecular basis of ion translocation in sodium/proton antiporters. Nat Struct Mol Biol 23: 248–255

8. Corey RA, Song W, Duncan AL, Ansell TB, Sansom MSP, Stansfeld PJ (2021) Identification and assessment of cardiolipin interactions with E. coli inner membrane proteins. Sci Adv 7

9. Daniel Asarnow EP, & Yifan Cheng., 2019. UCSF pyem v0.5. Zenodo. p. 10.5281/zenodo.3576630.

10. de Lartigue J, Polson H, Feldman M, Shokat K, Tooze SA, Urbe S, Clague MJ (2009) PIKfyve regulation of endosome-linked pathways. Traffic 10: 883–893

11. Dong Y, Gao Y, Ilie A, Kim D, Boucher A, Li B, Zhang XC, Orlowski J, Zhao Y (2021) Structure and mechanism of the human NHE1-CHP1 complex. Nat Commun 12: 3474

12. Dong Y, Li H, Ilie A, Gao Y, Boucher A, Zhang XC, Orlowski J, Zhao Y (2022) Structural basis of autoinhibition of the human NHE3-CHP1 complex. Sci Adv 8: eabn3925

13. Donowitz M, Ming Tse C, Fuster D (2013) SLC9/NHE gene family, a plasma membrane and organellar family of Na(+)/H(+) exchangers. Mol Aspects Med 34: 236–251

14. Donowitz M, Mohan S, Zhu CX, Chen TE, Lin R, Cha B, Zachos NC, Murtazina R, Sarker R, Li X (2009) NHE3 regulatory complexes. J Exp Biol 212: 1638–1646

15. Drew D, Boudker O (2016) Shared Molecular Mechanisms of Membrane Transporters. Annu Rev Biochem 85: 543–572

16. Drew D, Newstead S, Sonoda Y, Kim H, von Heijne G, Iwata S (2008) GFP-based optimization scheme for the overexpression and purification of eukaryotic membrane proteins in Saccharomyces cerevisiae. Nat Protoc 3: 784–798

17. Emsley P, Lohkamp B, Scott WG, Cowtan K (2010) Features and development of Coot. Acta Crystallogr D Biol Crystallogr 66: 486–501

18. Fliegel L (2019) Structural and Functional Changes in the Na(+)/H(+) Exchanger Isoform 1, Induced by Erk1/2 Phosphorylation. Int J Mol Sci 20

19. Forrest LR (2015) Structural Symmetry in Membrane Proteins. Annu Rev Biophys 44: 311–337

20. Fuster DG, Alexander RT (2014) Traditional and emerging roles for the SLC9 Na+/H+ exchangers. Pflugers Archiv: European journal of physiology 466: 61–76

21. Gupta K, Donlan JAC, Hopper JTS, Uzdavinys P, Landreh M, Struwe WB, Drew D, Baldwin AJ, Stansfeld PJ, Robinson CV (2017) The role of interfacial lipids in stabilizing membrane protein oligomers. Nature 541: 421–424

22. Hasegawa J, Strunk BS, Weisman LS (2017) PI5P and PI(3,5)P2: Minor, but Essential Phosphoinositides. Cell Struct Funct 42: 49–60

23. Hattori M, Hibbs RE, Gouaux E (2012) A fluorescence-detection size-exclusion chromatography-based thermostability assay for membrane protein precrystallization screening. Structure 20: 1293–1299

24. Ho CY, Alghamdi TA, Botelho RJ (2012) Phosphatidylinositol-3,5-bisphosphate: no longer the poor PIP2. Traffic 13: 1–8

25. Hunte C, Screpanti E, Venturi M, Rimon A, Padan E, Michel H (2005a) Structure of a Na+/H+ antiporter and insights into mechanism of action and regulation by pH. Nature 435: 1197–1202

26. Hunte C, Screpanti E, Venturi M, Rimon A, Padan E, Michel H (2005b) Structure of a Na+/H+ antiporter and insights into mechanism of action and regulation by pH. Nature 435: 1197–1202

27. Jumper J, Evans R, Pritzel A, Green T, Figurnov M, Ronneberger O, Tunyasuvunakool K, Bates R, Zidek A, Potapenko A et al (2021) Highly accurate protein structure prediction with AlphaFold. Nature 596: 583–589

28. Kondapalli KC, Llongueras JP, Capilla-Gonzalez V, Prasad H, Hack A, Smith C, Guerrero-Cazares H, Quinones-Hinojosa A, Rao R (2015) A leak pathway for luminal protons in endosomes drives oncogenic signalling in glioblastoma. Nat Commun 6: 6289

29. Kondapalli KC, Prasad H, Rao R (2014) An inside job: how endosomal Na(+)/H(+) exchangers link to autism and neurological disease. Front Cell Neurosci 8: 172

30. Krecek P, Skupa P, Libus J, Naramoto S, Tejos R, Friml J, Zazimalova E (2009) The PIN-FORMED (PIN) protein family of auxin transporters. Genome Biol 10: 249

31. Krulwich TA, Sachs G, Padan E (2011) Molecular aspects of bacterial pH sensing and homeostasis. Nature reviews Microbiology 9: 330–343

32. Lacroix J, Poet M, Maehrel C, Counillon L (2004) A mechanism for the activation of the Na/H exchanger NHE-1 by cytoplasmic acidification and mitogens. EMBO Rep 5: 91–96

33. Landreh M, Marklund EG, Uzdavinys P, Degiacomi MT, Coincon M, Gault J, Gupta K, Liko I, Benesch JL, Drew D et al (2017) Integrating mass spectrometry with MD simulations reveals the role of lipids in Na(+)/H(+) antiporters. Nature communications 8: 13993

34. Lee C, Kang HJ, von Ballmoos C, Newstead S, Uzdavinys P, Dotson DL, Iwata S, Beckstein O, Cameron AD, Drew D (2013) A two-domain elevator mechanism for sodium/proton antiport. Nature 501: 573–577

35. Lee C, Yashiro S, Dotson DL, Uzdavinys P, Iwata S, Sansom MS, von Ballmoos C, Beckstein O, Drew D, Cameron AD (2014) Crystal structure of the sodium-proton antiporter NhaA dimer and new mechanistic insights. The Journal of general physiology 144: 529–544

36. Li SC, Diakov TT, Xu T, Tarsio M, Zhu W, Couoh-Cardel S, Weisman LS, Kane PM (2014) The signaling lipid PI(3,5)P(2) stabilizes V(1)-V(o) sector interactions and activates the V-ATPase. Mol Biol Cell 25: 1251–1262

37. Masrati G, Dwivedi M, Rimon A, Gluck-Margolin Y, Kessel A, Ashkenazy H, Mayrose I, Padan E, Ben-Tal N (2018) Broad phylogenetic analysis of cation/proton antiporters reveals transport determinants. Nat Commun 9: 4205

38. Matsuoka R, Fudim R, Jung S, Zhang C, Bazzone A, Chatzikyriakidou Y, Robinson CV, Nomura N, Iwata S, Landreh M et al (2022) Structure, mechanism and lipid-mediated remodeling of the mammalian Na(+)/H(+) exchanger NHA2. Nat Struct Mol Biol 29: 108–120

39. Milosavljevic N, Monet M, Lena I, Brau F, Lacas-Gervais S, Feliciangeli S, Counillon L, Poet M (2014) The intracellular Na(+)/H(+) exchanger NHE7 effects a Na(+)-coupled, but not K(+)-coupled proton-loading mechanism in endocytosis. Cell Rep 7: 689–696

40. Mravec J, Skupa P, Bailly A, Hoyerova K, Krecek P, Bielach A, Petrasek J, Zhang J, Gaykova V, Stierhof YD et al (2009) Subcellular homeostasis of phytohormone auxin is mediated by the ER-localized PIN5 transporter. Nature 459: 1136–1140

41. Nakamura N, Tanaka S, Teko Y, Mitsui K, Kanazawa H (2005) Four Na+/H+ exchanger isoforms are distributed to Golgi and post-Golgi compartments and are involved in organelle pH regulation. J Biol Chem 280: 1561–1572

42. Nji E, Chatzikyriakidou Y, Landreh M, Drew D (2018) An engineered thermal-shift screen reveals specific lipid preferences of eukaryotic and prokaryotic membrane proteins. Nat Commun 9: 4253

43. Norholm AB, Hendus-Altenburger R, Bjerre G, Kjaergaard M, Pedersen SF, Kragelund BB (2011) The intracellular distal tail of the Na+/H+ exchanger NHE1 is intrinsically disordered: implications for NHE1 trafficking. Biochemistry 50: 3469–3480

44. Odunewu-Aderibigbe A, Fliegel L (2014) The Na(+) /H(+) exchanger and pH regulation in the heart. IUBMB Life 66: 679–685

45. Okazaki KI, Wohlert D, Warnau J, Jung H, Yildiz O, Kuhlbrandt W, Hummer G (2019) Mechanism of the electroneutral sodium/proton antiporter PaNhaP from transition-path shooting. Nat Commun 10: 1742

46. Orlowski J, Grinstein S (2004) Diversity of the mammalian sodium/proton exchanger SLC9 gene family. Pflugers Archiv: European journal of physiology 447: 549–565

47. Padan E (2008) The enlightening encounter between structure and function in the NhaA Na+-H+ antiporter. Trends Biochem Sci 33: 435–443

48. Padan E, Tzubery T, Herz K, Kozachkov L, Rimon A, Galili L (2004) NhaA of Escherichia coli, as a model of a pH-regulated Na+/H+antiporter. Biochimica et biophysica acta 1658: 2–13

49. Paulino C, Wohlert D, Kapotova E, Yildiz O, Kuhlbrandt W (2014) Structure and transport mechanism of the sodium/proton antiporter MjNhaP1. eLife 3: e03583

50. Pedersen SF, Counillon L (2019) The SLC9A-C Mammalian Na(+)/H(+) Exchanger Family: Molecules, Mechanisms, and Physiology. Physiol Rev 99: 2015–2113

51. Pettersen EF, Goddard TD, Huang CC, Couch GS, Greenblatt DM, Meng EC, Ferrin TE (2004) UCSF Chimera--a visualization system for exploratory research and analysis. J Comput Chem 25: 1605–1612

52. Pettersen EF, Goddard TD, Huang CC, Meng EC, Couch GS, Croll TI, Morris JH, Ferrin TE (2021) UCSF ChimeraX: Structure visualization for researchers, educators, and developers. Protein Sci 30: 70–82

53. Pouyssegur J, Sardet C, Franchi A, L’Allemain G, Paris S (1984) A specific mutation abolishing Na+/H+ antiport activity in hamster fibroblasts precludes growth at neutral and acidic pH. Proc Natl Acad Sci U S A 81: 4833–4837

54. Punjani A, Rubinstein JL, Fleet DJ, Brubaker MA (2017) cryoSPARC: algorithms for rapid unsupervised cryo-EM structure determination. Nat Methods 14: 290–296

55. Qi Y, Ingolfsson HI, Cheng X, Lee J, Marrink SJ, Im W (2015) CHARMM-GUI Martini Maker for Coarse-Grained Simulations with the Martini Force Field. J Chem Theory Comput 11: 4486–4494

56. Quick M, Dwivedi M, Padan E (2021) Insight into the direct interaction of Na(+) with NhaA and mechanistic implications. Sci Rep 11: 7045

57. Rimon A, Mondal R, Friedler A, Padan E (2019) Cardiolipin is an Optimal Phospholipid for the Assembly, Stability, and Proper Functionality of the Dimeric Form of NhaA Na(+)/H(+) Antiporter. Sci Rep 9: 17662

58. Rohou A, Grigorieff N (2015) CTFFIND4: Fast and accurate defocus estimation from electron micrographs. J Struct Biol 192: 216–221

59. Romantsov T, Guan Z, Wood JM (2009) Cardiolipin and the osmotic stress responses of bacteria. Biochim Biophys Acta 1788: 2092–2100

60. Simonin A, Fuster D (2010) Nedd4-1 and beta-arrestin-1 are key regulators of Na+/H+ exchanger 1 ubiquitylation, endocytosis, and function. J Biol Chem 285: 38293–38303

61. Sjogaard-Frich LM, Prestel A, Pedersen ES, Severin M, Kristensen KK, Olsen JG, Kragelund BB, Pedersen SF (2021) Dynamic Na(+)/H(+) exchanger 1 (NHE1) – calmodulin complexes of varying stoichiometry and structure regulate Ca(2+)-dependent NHE1 activation. eLife 10

62. Slepkov ER, Rainey JK, Sykes BD, Fliegel L (2007) Structural and functional analysis of the Na+/H+ exchanger. Biochem J 401: 623–633

63. Su N, Zhu A, Tao X, Ding ZJ, Chang S, Ye F, Zhang Y, Zhao C, Chen Q, Wang J et al (2022) Structures and mechanisms of the Arabidopsis auxin transporter PIN3. Nature

64. Suades A, Qureshi A, McComas SE, Coincon M, Rudling A, Chatzikyriakidou Y, Landreh M, Carlsson J, Drew D (2023) Establishing mammalian GLUT kinetics and lipid composition influences in a reconstituted-liposome system. Nat Commun 14: 4070

65. Teale WD, Pasternak T, Dal Bosco C, Dovzhenko A, Kratzat K, Bildl W, Schworer M, Falk T, Ruperti B, Schaefer JV et al (2021) Flavonol-mediated stabilization of PIN efflux complexes regulates polar auxin transport. The EMBO journal 40: e104416

66. Ung KL, Winkler M, Schulz L, Kolb M, Janacek DP, Dedic E, Stokes DL, Hammes UZ, Pedersen BP (2022) Structures and mechanism of the plant PIN-FORMED auxin transporter. Nature

67. Wakabayashi S, Fafournoux P, Sardet C, Pouyssegur J (1992) The Na+/H+ antiporter cytoplasmic domain mediates growth factor signals and controls “H(+)-sensing”. Proc Natl Acad Sci U S A 89: 2424–2428

68. Wakabayashi S, Ikeda T, Iwamoto T, Pouyssegur J, Shigekawa M (1997) Calmodulin-binding autoinhibitory domain controls “pH-sensing” in the Na+/H+ exchanger NHE1 through sequence-specific interaction. Biochemistry 36: 12854–12861

69. Winkelmann I, Uzdavinys P, Kenney IM, Brock J, Meier PF, Wagner LM, Gabriel F, Jung S, Matsuoka R, von Ballmoos C et al (2022) Crystal structure of the Na(+)/H(+) antiporter NhaA at active pH reveals the mechanistic basis for pH sensing. Nat Commun 13: 6383

70. Winklemann I, Matsuoka R, Meier PF, Shutin D, Zhang C, Orellana L, Sexton R, Landreh M, Robinson CV, Beckstein O et al (2020) Structure and elevator mechanism of the mammalian sodium/proton exchanger NHE9. The EMBO journal 39: e105908

71. Wohlert D, Kuhlbrandt W, Yildiz O (2014) Structure and substrate ion binding in the sodium/proton antiporter PaNhaP. eLife 3: e03579

72. Yang Z, Xia J, Hong J, Zhang C, Wei H, Ying W, Sun C, Sun L, Mao Y, Gao Y et al (2022) Structural insights into auxin recognition and efflux by Arabidopsis PIN1. Nature

73. Yeo H, Mehta V, Gulati A, Drew D (2023) Structure and electromechanical coupling of a voltage-gated Na(+)/H(+) exchanger. Nature 623: 193–201

74. Zhang-James Y, Vaudel M, Mjaavatten O, Berven FS, Haavik J, Faraone SV (2019) Effect of disease-associated SLC9A9 mutations on protein-protein interaction networks: implications for molecular mechanisms for ADHD and autism. Atten Defic Hyperact Disord 11: 91–105

75. Zheng SQ, Palovcak E, Armache JP, Verba KA, Cheng Y, Agard DA (2017) MotionCor2: anisotropic correction of beam-induced motion for improved cryo-electron microscopy. Nat Methods 14: 331–332

76. Zivanov J, Nakane T, Forsberg BO, Kimanius D, Hagen WJ, Lindahl E, Scheres SH (2018) New tools for automated high-resolution cryo-EM structure determination in RELION-3. eLife 7

