## Supplementary Figures and Table 1 for "PI-(3,5)P2-mediated oligomerization of the endosomal sodium/proton exchanger NHE9"

Supplementary Fig. 1

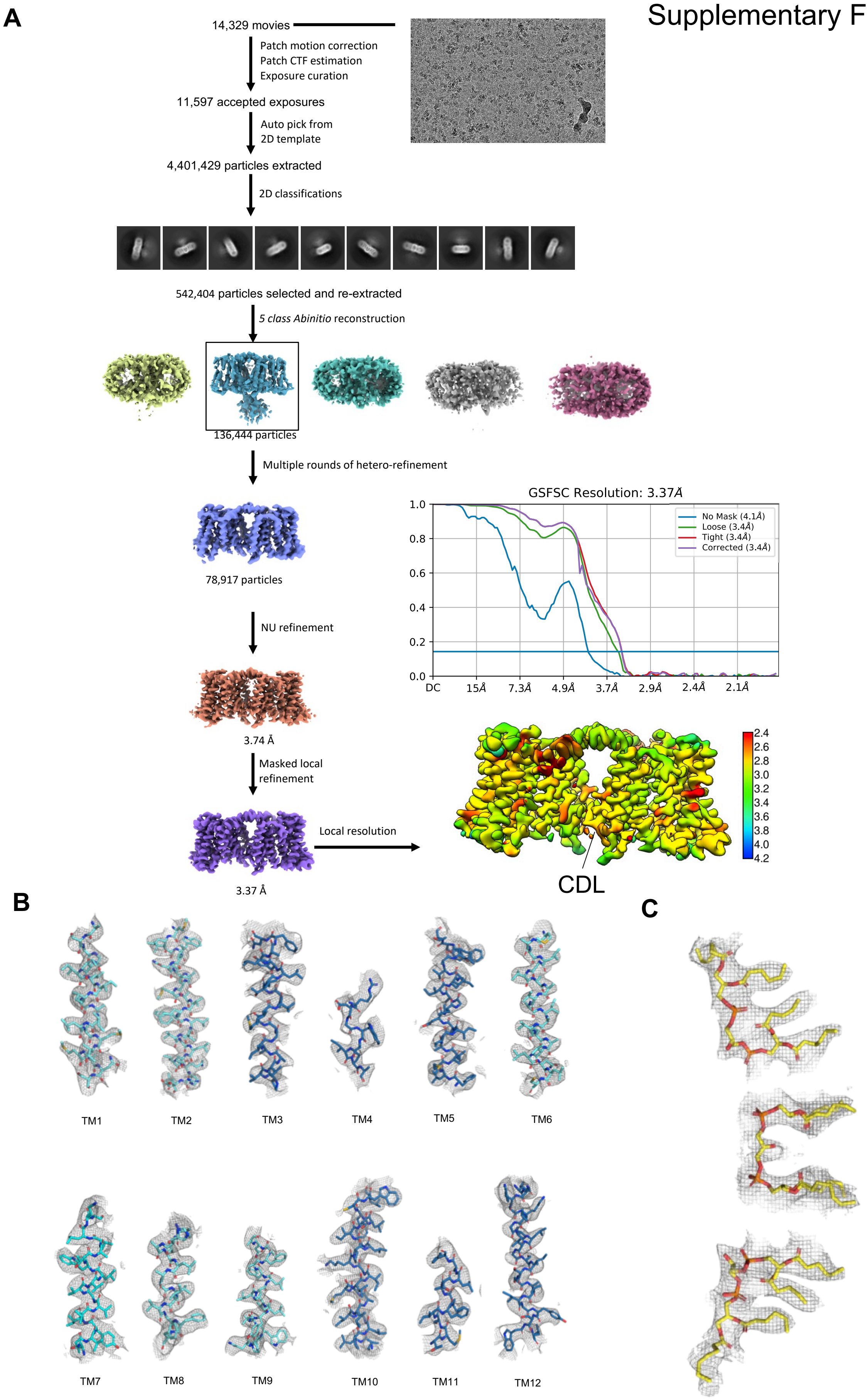

**Supplementary Fig. 1. The data-processing workflow of EcNhaAmut2 at pH 7.5. A.** The dataset contained 14,329 movies that were corrected by MotionCor2 and CTFFind. After reference-based auto-picking, 4,401,429 particles were picked. Several rounds of 2D classifications yielded in good 2D classes of 542,404 particles. 3D classification and 3D refinement of model resulted in an electron density map of 3.74 Å. The electron density map was subjected to micelle subtraction, local refinement, per particle CTF and Bayesian polishing.Final resolution of 3.37Å at gold-standard FSC (0.143), with a local resolution range of 2.4–3.6 Å. **B.** Cryo-EM density map and model are shown for all the transmembrane segments for NhaA with the dimer domain (cyan), transport domain (sky-blue), all shown as sticks. **C.** Cryo-EM density map and cardiolipin lipids (yellow-orange).

Supplementary Fig. 2

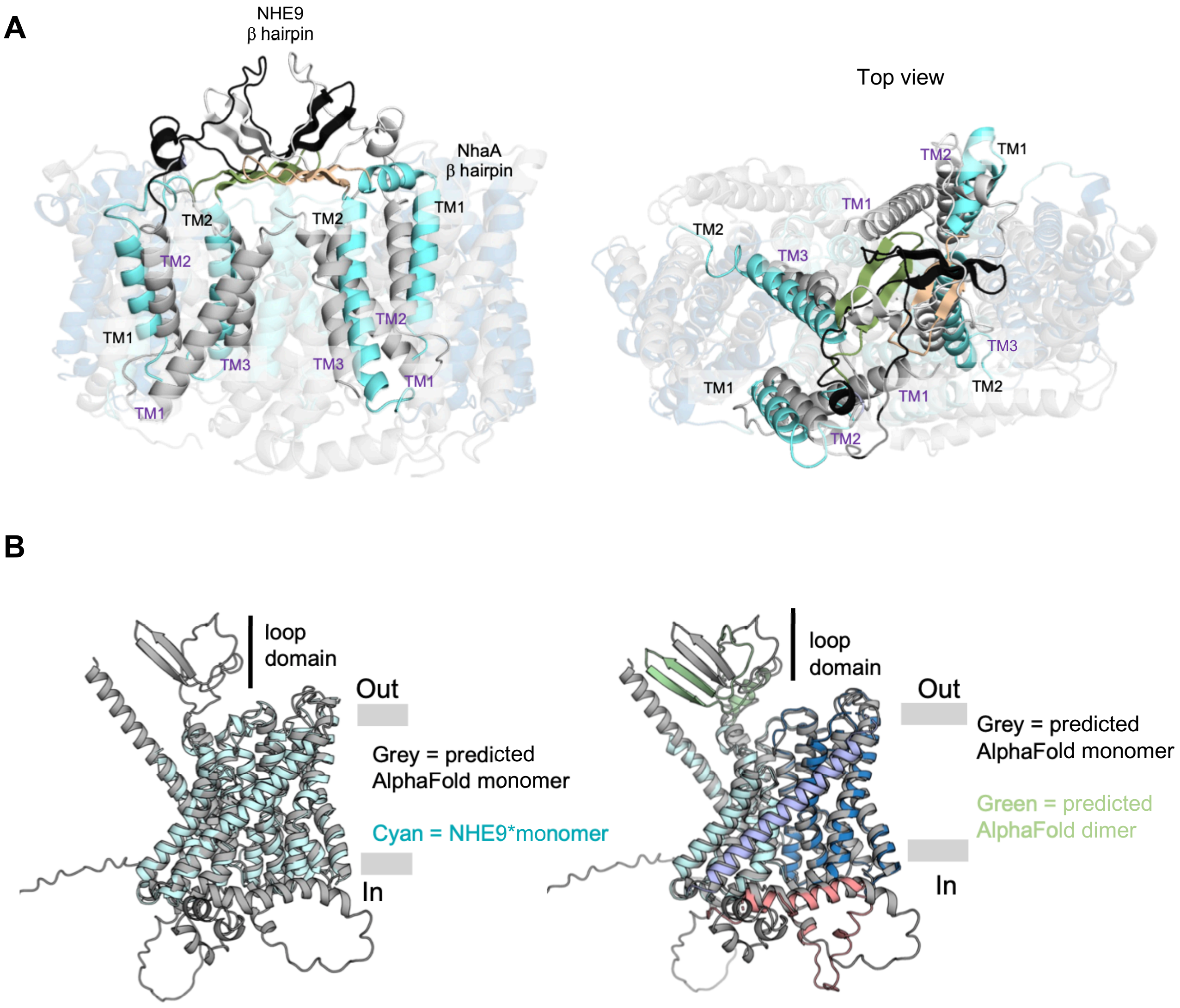


**Supplementary Figure 2. β-hairpin containing NhaA and NHE9 proteins. (A)** *left*: Superimposition of *Ec*NhaA dimer structure (coloured) and NHE9*CC dimer structure (grey) highlighting the topologically equivalent TM's connected to β-hairpin in both the structures. The β-hairpins of *Ec*NhaA (olive and wheat) are oriented parallel to membranes compared to NHE9*CC (black and grey) in vertically upright position. TM's of *Ec*NhaA (cyan) are marked in black whereas for NHE9*CC (grey) in purple. *right*: Topview of superimposed *Ec*NhaA dimer structure (coloured) and NHE9*CC dimer structure (grey) **(B)** *left*: Superimposition of the AlphaFold *horse* NHE9 monomer and *horse* NHE9 cryo EM structure (PDB: 6Z3Z). *right*: Superimposition of the AlphaFold *horse* NHE9 monomer and dimer models.

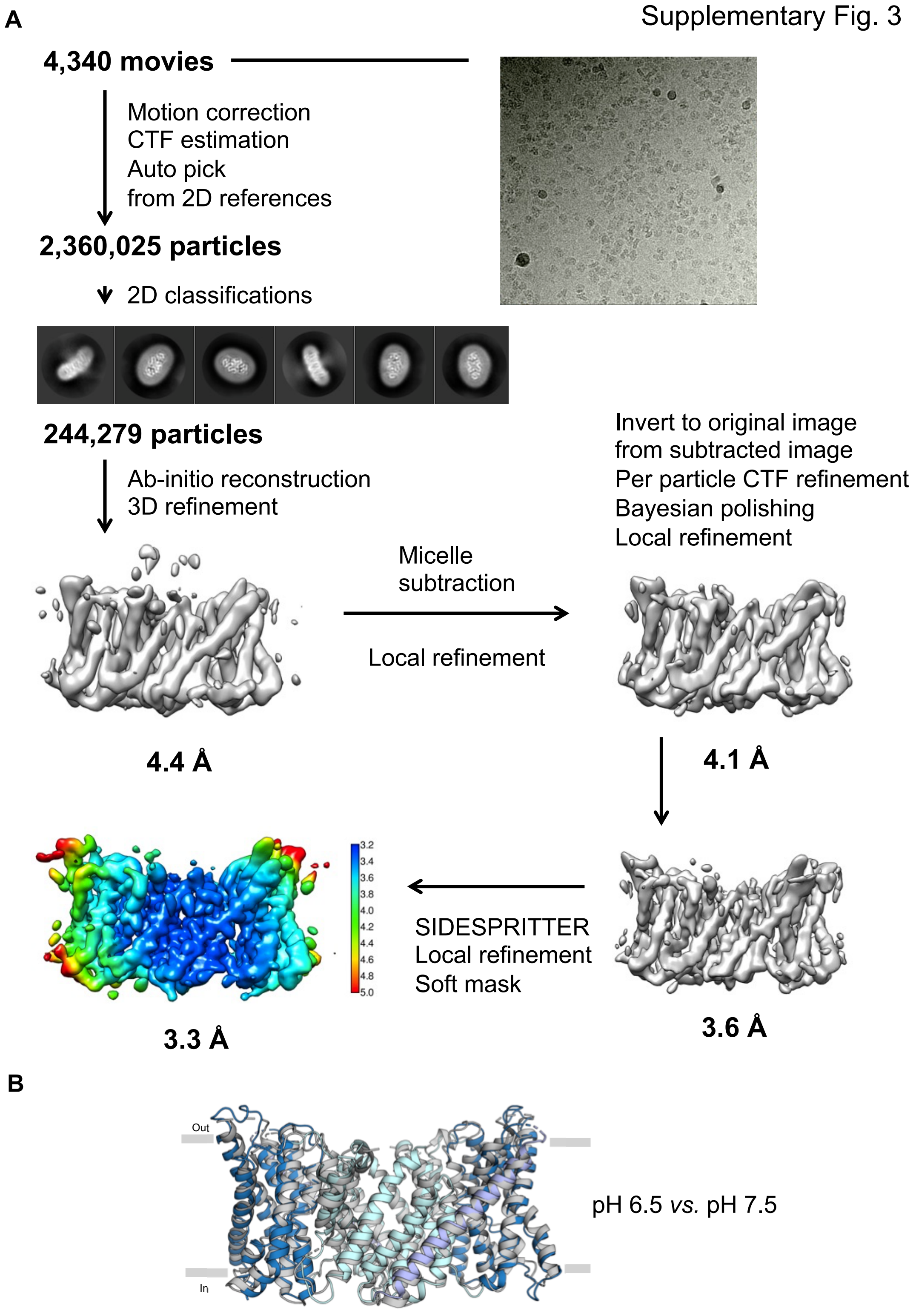


**Supplementary Figure 3. Data-processing workflow of *horse* NHE9*at pH 6.5.** The dataset contained 4,340 movies that were corrected by MotionCor2 and CTFFind. After reference-based auto-picking, 2,360,025 particles were picked. Several rounds of 2D classifications yielded in good 2D classes of 244,279 particles. 3D classification and 3D refinement of model resulted in an electron density map of 4.4 Å. The electron density map was subjected to micelle subtraction, local refinement, per particle CTF and Bayesian polishing. A final resolution of 3.31Å was achieved at gold-standard FSC (0.143), with a local resolution range of 3.2–5.0 Å. **B**. Superimposed NHE9* structure at pH 7.5 (grey; PDB ID: 6Z3Z) and NHE9* at pH 6.5 with dimerization domain (cyan), core domain (blue), loop domains (pale-green, sand), and TM7 linking helix (light-purple); the β-hairpin TM2-TM3 loop domain in the later modelled NHE9* structure at pH 6.5 is not shown here as it was not visible after masked refinement.

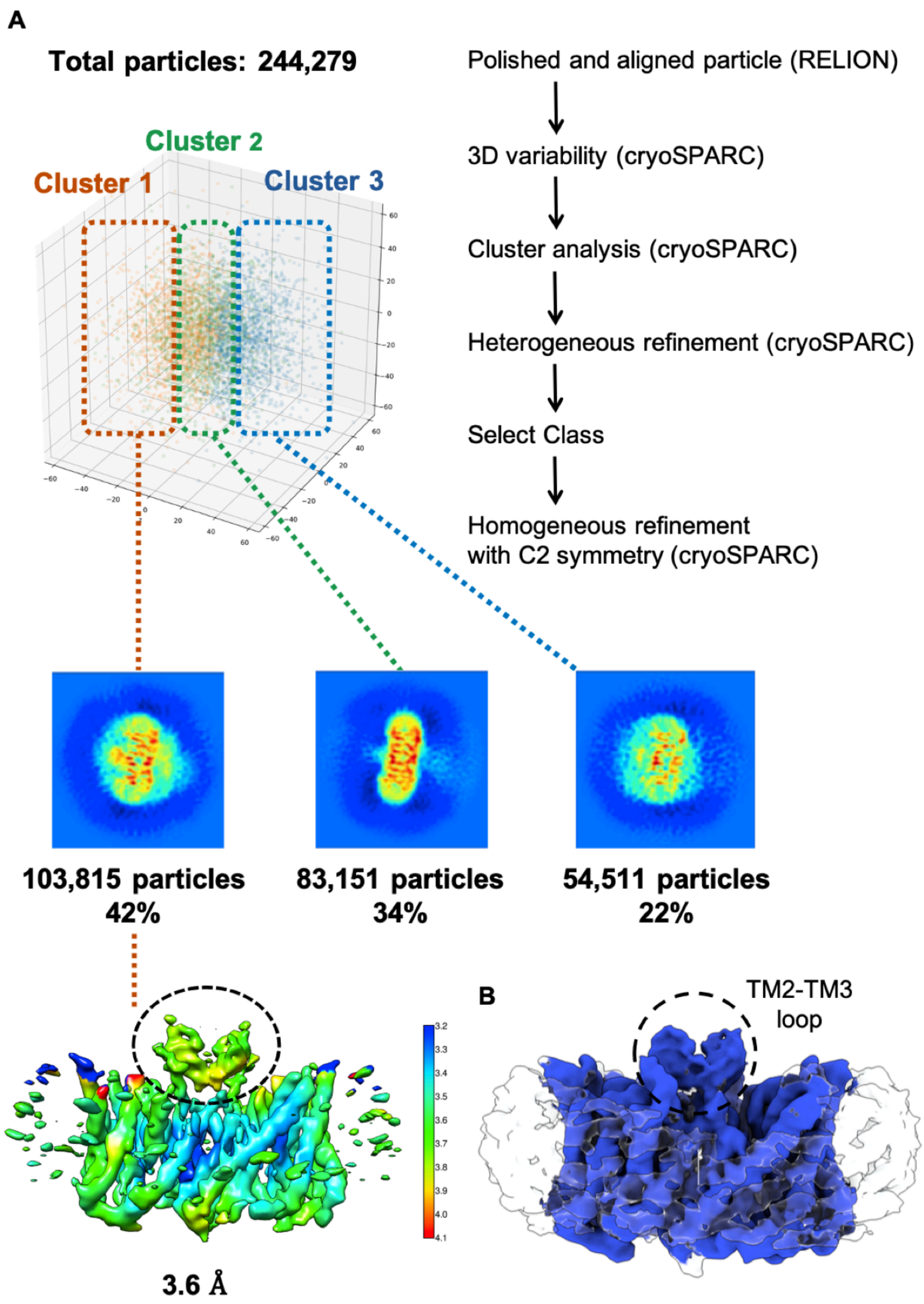


**Supplementary Figure 4. 3D Variability Analysis improved the overall resolution and the density for the TM2-TM3 β-hairpin loop domains. (A)** All particles of the final 3D reconstruction were subjected to 3D variability analysis followed by cluster analysis. Cluster 1 including 42% of particles were subsequently subjected to heterogeneous refinement followed by homogeneous refinement with C2 symmetry applied. A final resolution of 3.6Å was achieved at gold-standard FSC (0.143), with a local resolution range of 3.2–4.1 Å. Improved electron density for the loop domains above the dimerization domain (black dashed line) was observed. **(B)** Composite cryo- EM map combining the map features seen with masked refinement (Supplementary Fig. 3) and after 3D Variability and cluster analysis shown in (A).

A

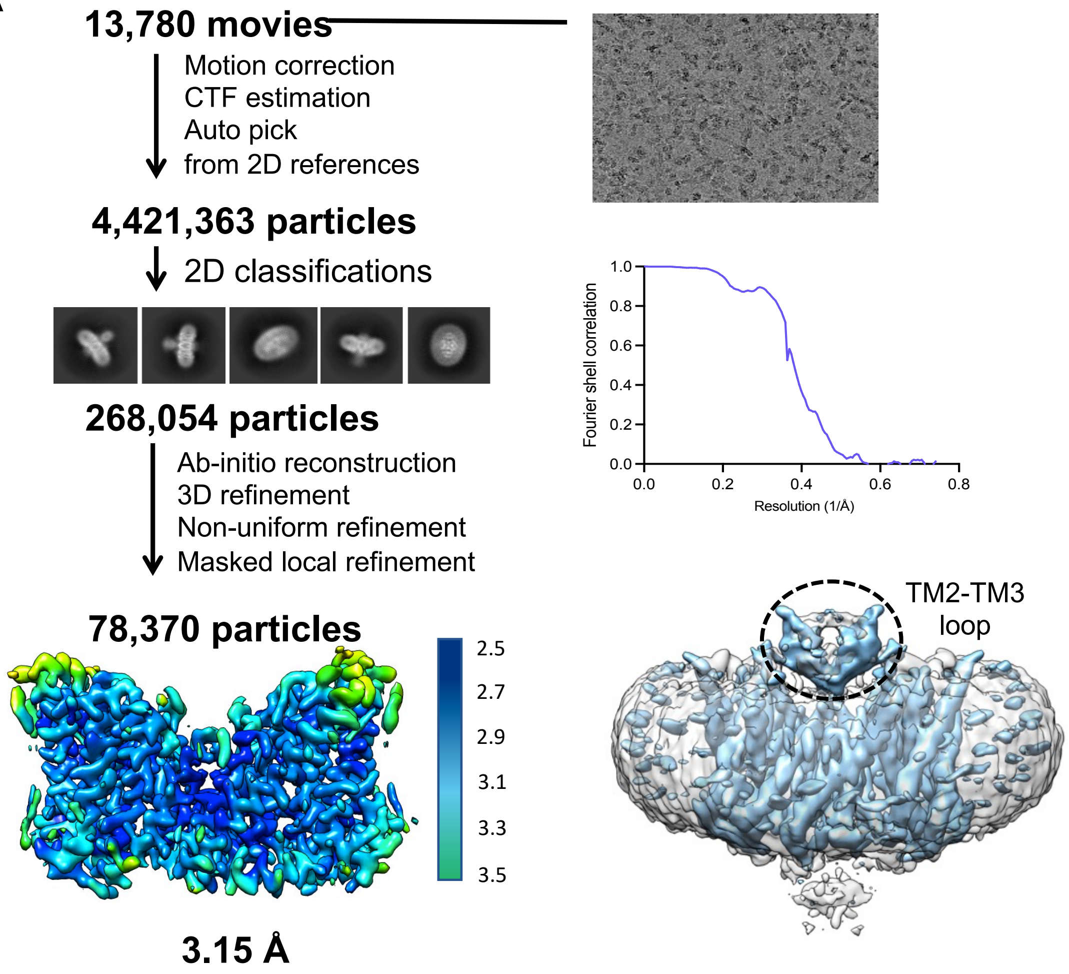


B

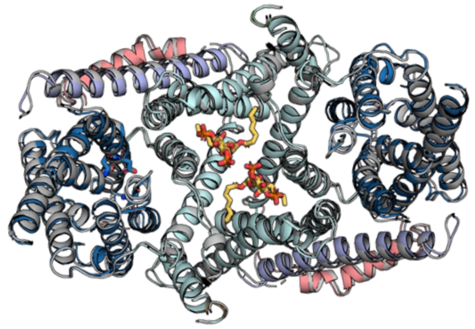

**Supplementary Figure 5. Cryo EM workflow and NHE9*CC structure. (A)** The dataset contained 13,780 movies that were corrected by MotionCor2 and CTFFind. After reference-based auto-picking, 4,421,363 particles were picked. Several rounds of 2D classifications yielded in good 2D classes of 133,145 particles. 3D classification, 3D refinement of the electron density map followed by micelle subtraction and local refinement resulted in a final resolution of 3.15 Å at gold-standard FSC (0.143). **(B)** Superimposed cryo-EM strucure of NHE9*loop (grey) and a subsequent variant NHE9*CC with core domain (cyan), transporter domain (blue), linker helix (light blue), CTD helix (salmon) and the $PI(3,5)P_2$ lipids as yellow sticks.

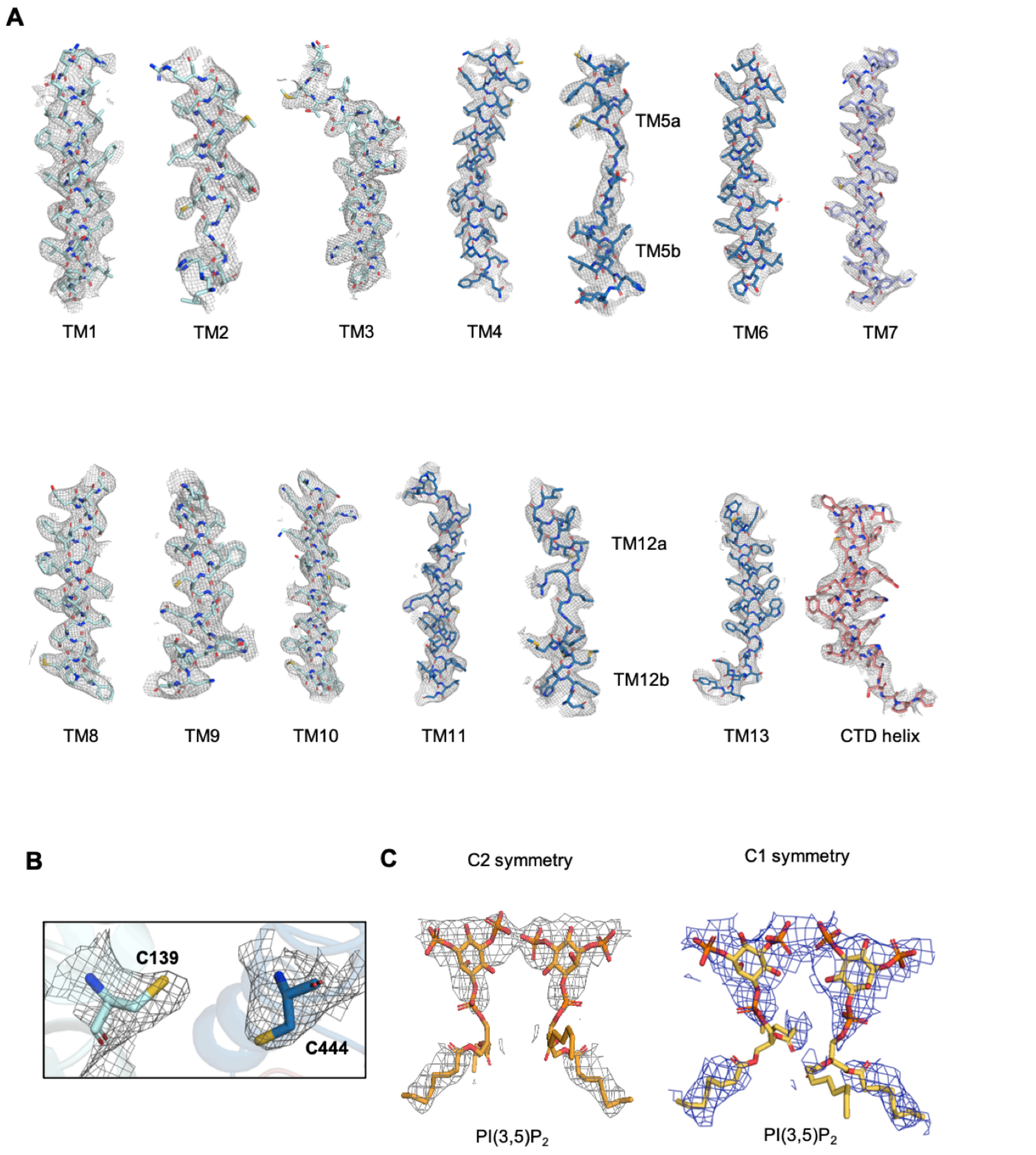


**Supplementary Figure 6. Cryo-EM desity of NHE9*CC. (A)** Cryo-EM desity map and model are shown for all the transmembrane segments for NHE9* in dimer domain (palecyan), transport domain (blue), and linker helix (lightpurple), PI(3,5)$P_2$ (yelloworange), helix of the CTD (salmon), all protein shown as sticks colored as in figure 1A. **(B)** Electron density map of the two mutated cysteine residues Leu139Cys and Ile444Cys does not show a disulphide bond. (**C**) Cryo-EM map density (blue mesh) around the lipid PI(3,5)$P_2$ (yellow sticks), with and without C2 symmetry applied during processing.

Supplementary Fig. 7

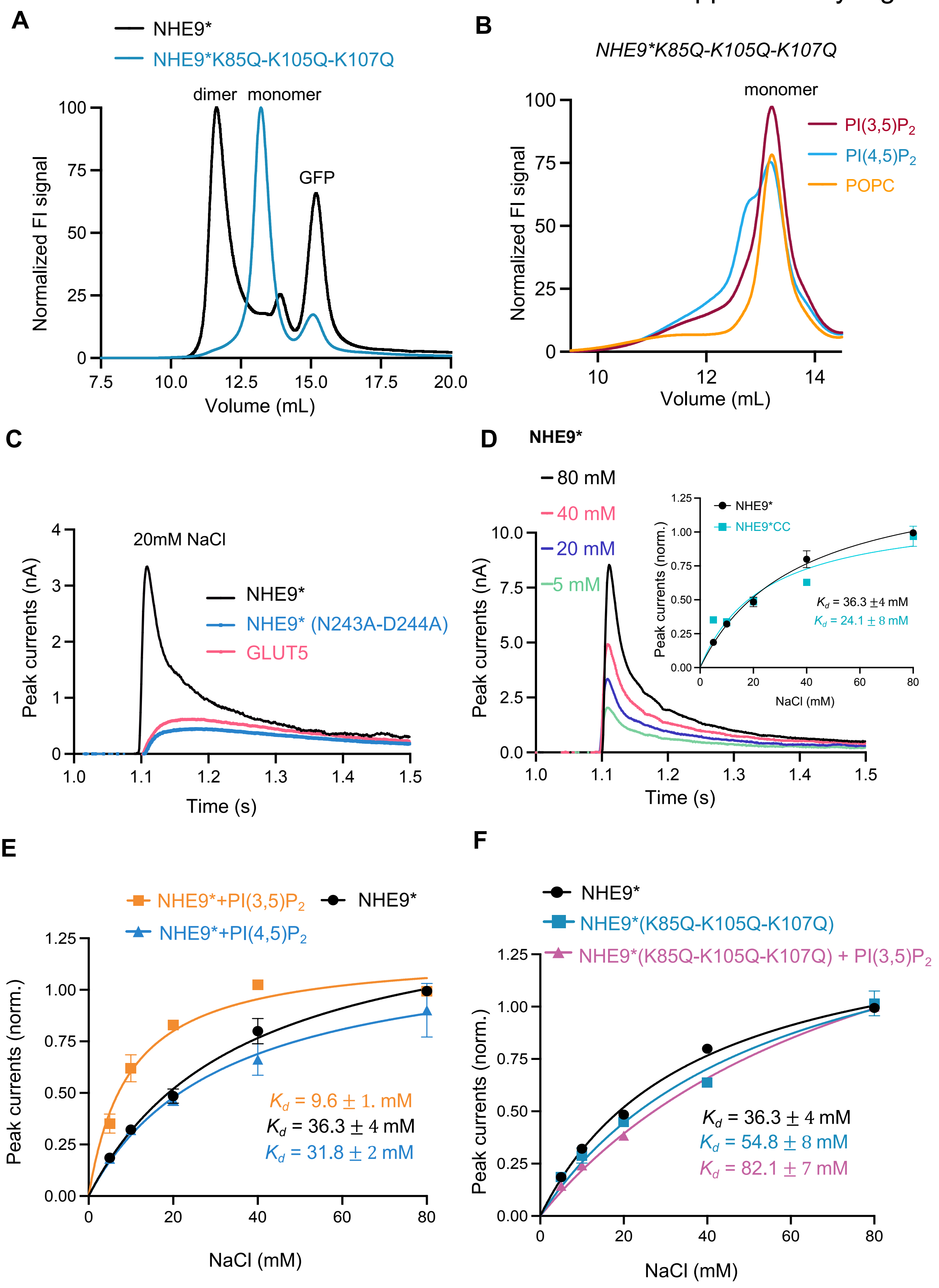

**Supplementary Fig. 7 NHE9* and varaints analysed by FSEC and SSM-based electrophysiology** (**A**). Representative normalized FSEC traces for purified NHE9*-GFP fusion (black) and NHE9*(K85Q, K105Q, K107Q)-GFP fusion (blue). (**B**) Representative normalized FSEC traces for purified NHE9*(K85Q, K105Q, K107Q)-GFP fusion after heating at 50ºC for 10 mins in the presence of DDM-solublised lipids as labelled. (**C**) Transient currents were recorded for NHE9* proteoliposomes (black trace) at symmetrical pH 7.5 after the addition of 20 mM NaCl. Peak currents for an ion-binding site NHE9* variant N243A-D244A (blue trace) and fructose transporter GLUT5 (Suades *et al*, 2023) (red trace) after the addition of 20 mM NaCl are also shown. (**D**) SSM-based electrophysiology measurements of NHE9* proteoliposomes with transient currents recorded after $Na^+$ concentration jumps at pH 7.5 on both sides. *Insert:* Fit of the normalized amplitude of the transient currents for NHE9* and NHE9*CC as a function of $Na^+$ concentrations and the corresponding binding affinity across n = 2 and 3 different sensors ($K_d$), respectivelty. Error bars are the SEM of n = 3 technical repeats (1 sensor) and the $K_d$ is the mean ± s.d. for n = 2 or 3 titrations (sensors). (**E**) Fit of the normalized amplitude of the transient currents for either NHE9* (black curve) or NHE9* pre-incubated with PI(4,5)P2 lipid (orange curve) or NHE9* pre-incubated with PI(3,5)P2 lipid (blue curve) as a function of $Na^+$ concentrations and the corresponding binding affinity across 3 different sensors ($K_d$). Error bars are the mean ± s.d. for n = 3 independent experiments (sensors). (**F**) Fit of the normalized amplitude of the transient currents for either NHE9* (black curve) or NHE9* β-hairpin lysine varaint (K85Q, K105Q, K107Q) pre-incubated with buffer (blue curve) or PI(3,5)P2 lipid (purple) as a function of $Na^+$ concentrations and the corresponding binding affinity across 3 different sensors ($K_d$). Error bars are the mean ± s.d. for n = 3 independent experiments (sensors).

Supplementary Fig. 8

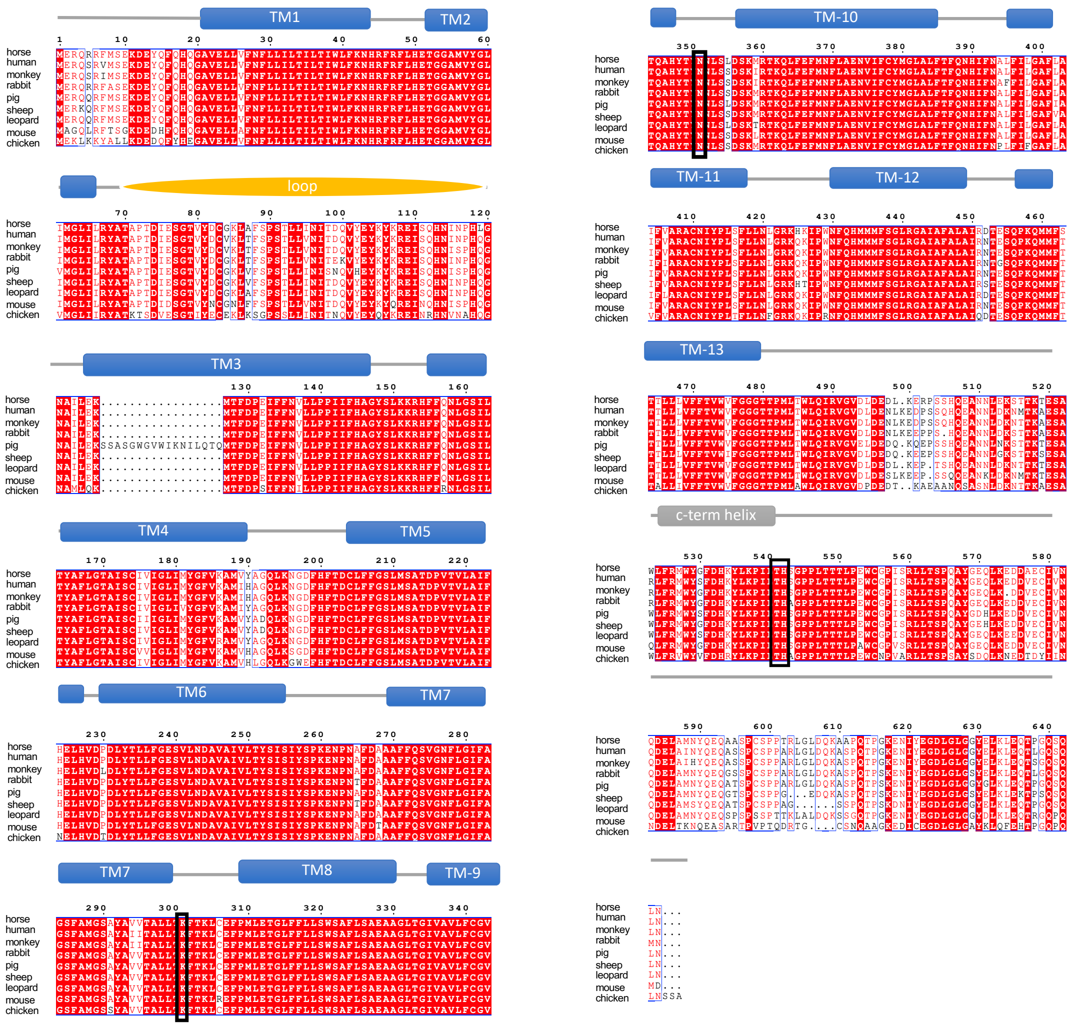

**Supplementary Figure 8. Multiple sequence alignment for NHE9 among vertebrates.** The following are the accession ID's for horse (F7B113), human(Q8IVB4), monkey (H9FT94), rabbit (G1SY59), pig(A0A287BEB2), sheep (W5P6Y8), leopard (A0A6P4V361), mouse (Q8BZ00), chicken (A0A8V0YVM6),. Residues with over 90% sequence identity are indicated by red background. Positions which are involved in gating (K301) and c-terminus regulation (T541, H542) are highlighted in black box. Residues which form TMs (blue), Loop (yellow) and C-term helix (grey) are indicated.

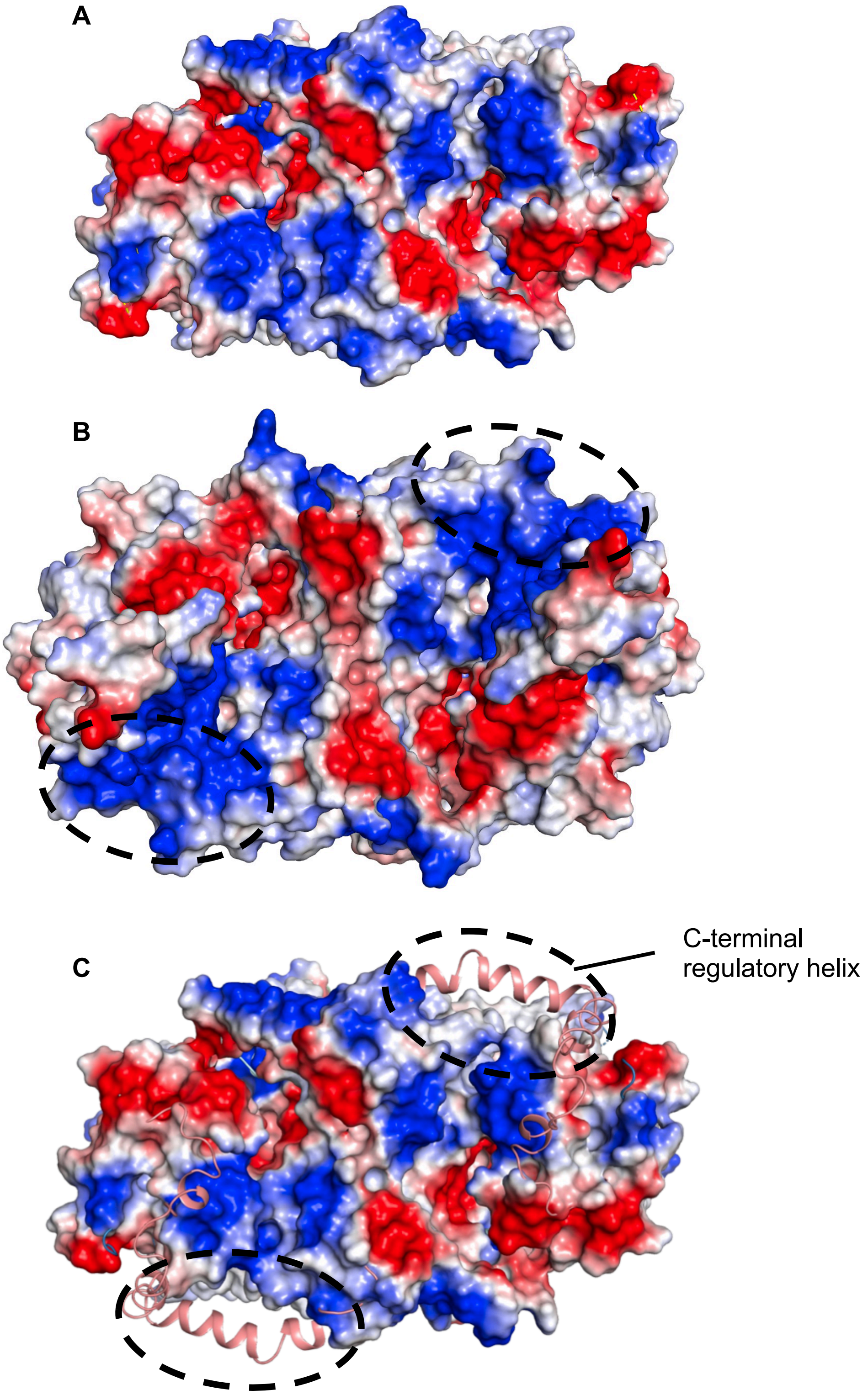


**Supplementary Figure 9. Visualizing the difference upon inclusion of the C-terminal regulatory domain on overall structure of NHE9 viewed from the cytosloic side. (A)** Previous published structure of NHE9* (PDB id: 6Z3Z) where the CTD could not be modelled shown in as an electrostatic surface potential. **(B)** Improved NHE9* CC structure with CTD density (black dashed circle) (PDB id; not deposited. **(C)** Improved NHE9* CC structure shown as in A with CTD helix built into the additional map density (black dashed circle).

| | NHE9*loop | NHE9*CC | NhaA-mut2 |
|---|---|---|---|
| | Data collection and processing statistics | | |
| Magnification | 165,000 | 130,000 | 130,000 |
| Voltage(kV) | 300 | 300 | 300 |
| Electron exposure ($e^-$/Å$^2$ ) | 57 | 68.5 | 68.11 |
| Defocus range (µm) | 0.9-2.3 | 0.4-2.0 | 0.6-2.0 |
| Pixel size (Å) | 0.82 | 0.66 | 0.6645 |
| Symmetry imposed | C2 | C2 (and C1) | C2 |
| Initial particle images (no.) | 2,360,025 | 4,421,363 | 4,401,429 |
| Final particle images (no.) | 103,815 | 78,370 | 78,917 |
| Map resolution (Å) | 3.6 | 3.15 | 3.37 |
| FSC threshold | 0.143 | 0.143 | 0.143 |
| Map resolution range | 3.2-4.1 | 2.5-3.5 | 2.4-4.0 |
| | Refinement | | |
| Initiial model used (PDB code) | NHE9*CC and NHE9 Alphafold2 dimer | 6Z3Z and NHE9 Alphafold2 dimer | 7S24 |
| Model resolution (Å) | 6.7 | 3.5 | 3.8 |
| FSC threshold | 0.5 | 0.5 | 0.5 |
| Model composition | | | |
| Non-hydrogen atoms | 8420 | 8506 | 5739 |
| Protein residues | 1058 | 1058 | 744 |
| Ligands | - | PI(3,5)$P_2$ | CDL:3 |
| B factor (Å$^2$) | | | |
| Protein | 237.07 | 165.98 | 92.07 |
| Ligand | - | 221.06 | 161.24 |
| R.m.s deviations | | | |
| Bond lengths (Å) | 0.001 | 0.002 | 0.002 |
| Bond angles (°) | 0.394 | 0.508 | 0.638 |
| Validation | | | |
| Mol probity score | 1.56 | 1.80 | 1.73 |
| Clash score | 4.82 | 7.95 | 6.59 |
| Poor rotamers(%) | 1.11 | 0.00 | 0.86 |
| Ramachandran plot | | | |
| Favoured (%) | 96.0 | 94.65 | 94.58 |
| Allowed (%) | 4.0 | 5.35 | 5.42 |
| Disallowed (%) | 0 | 0 | 0 |

**Supplementary Table 1. Data collection, processing and refinement statistics of NHE9*, NHE9*CC and *Ec*NhaA-mut2 cryo EM structures.**
